# Rewiring *Saccharomyces cerevisiae* metabolism for optimised Taxol® precursors production

**DOI:** 10.1101/2023.06.03.543533

**Authors:** Behnaz Nowrouzi, Pablo Torres-Montero, Eduard J. Kerkhoven, José L. Martínez, Leonardo Rios-Solis

**Affiliations:** Institute for Bioengineering, School of Engineering, The University of Edinburgh, Edinburgh, EH9 3BF United Kingdom; Centre for Engineering Biology, The University of Edinburgh, Edinburgh, EH9 3BD United Kingdom; School of Natural and Environmental Sciences, Molecular Biology and Biotechnology Division, Newcastle University, Newcastle upon Tyne NE1 7RU United Kingdom; Department of Biology and Biological Engineering, Chalmers University of Technology, SE-412 96 Gothenburg, Sweden; Novo Nordisk Foundation Center for Biosustainability, Chalmers University of Technology, SE-412 96 Gothenburg, Sweden; Department of Biotechnology and Biomedicine, Technical University of Denmark, Søltofts Plads Building 223, Kgs. Lyngby, 2800, Denmark; Department of Biochemical Engineering, The Advanced Centre for Biochemical Engineering, University College London, Gower Street, London, WC1E 6BT, UK

**Keywords:** *Saccharomyces cerevisiae*, Taxol®, Taxadiene, Oxidative Stress, Adaptive Laboratory Evolution

## Abstract

*Saccharomyces cerevisiae* has been recognised as a convenient host for the production of early precursors to the Taxol® anticancer drug. Recent studies have highlighted the harmful impact of oxidative stress as a result of the activity of Taxol® first cytochrome P450-reductase enzymes (*Taxus* spp. CYP725A4-POR). Here, we evolved a new oxidative stress-tolerant yeast strain on galactose, which led to a three-fold higher titre of the CYP725A4 enzyme substrate, taxadiene. We comprehensively analysed the performance of the evolved and parent strain in galactose-limited chemostat cultures before and during oxidative stress induction. Integrating the transcriptomics and metabolite profiling data in an enzyme-constrained genome scale model enabled a more accurate prediction of changes that occurred to biological pathways as a response to/consequence of evolution and oxidative stress. The analyses showed a better performance of the evolved strain with improved respiration and reduced overflow metabolites production. The strain was robust to re-introduction of the oxidative stress, potentially due to the cross-protection mechanism, which contributed to likely better heme, flavin and NADPH availability for an optimal expression of *CYP725A4* and *POR* in yeast. The increased level of taxadiene production has potentially occurred due to the antioxidant properties of taxadiene or as a mechanism to overcome the toxicity of geranylgeranyl diphosphate, the precursor to taxadiene synthase.

**Highlights:**

- The antioxidant properties of taxadiene promotes its production in *Saccharomyces cerevisiae*
- *S. cerevisiae* ALE on H_2_O_2_ and galactose regulates Flavin, iron and NADPH metabolism as well as carbon and protein recycling pathways through cross-protection and anticipation mechanisms

Figure 1.
Graphical abstract of the study.
Figure was created with BioRender.com.

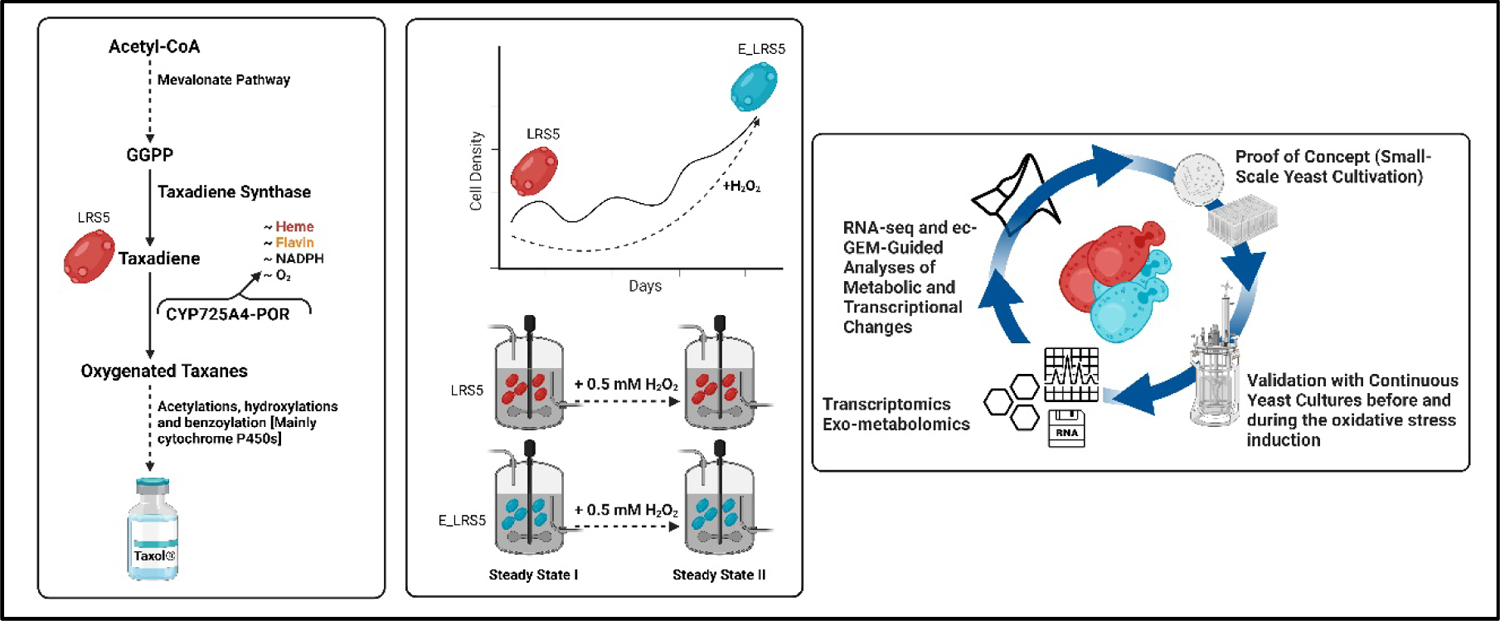

## 1. Introduction

Originally extracted from *Taxus* spp., Taxol® (paclitaxel) is one of the world’s best-selling anticancer drugs. While its full biosynthesis pathway has not been elucidated yet, certainly the dominance of cytochrome P450 enzymatic complexes with multiple domains and cofactor requirements including flavins, NADPH and irons (Nowrouzi et al., 2022) has hindered sufficient progress in total drug synthesis by heterologous systems. As a result, the main attention has so far been on optimising the early enzymatic steps. Briefly, Taxol® biosynthesis commences with the conversion of geranylgeranyl diphosphate (GGPP) into the first taxane backbone, taxa-4(5),11(12)-diene (taxadiene) and its isomer, taxa-4(20),11(12)-diene (iso-taxadiene), followed by their hydroxylation by the first cytochrome P450, CYP725A4 enzyme, to yield the first diterpenoid/oxygenated taxane, taxadiene-5α-ol (T5α-ol), together with other side diterpenoids (Barton et al., 2016; Edgar et al., 2016; Nowrouzi et al., 2022, Walls et al., 2022b). From there, an array of acetylation, hydroxylation and benzoylation furnishing events lead to Taxol® formation (Lange and Conner, 2021; Walker and Croteau, 2000).

We have previously reported on the construction of a high-taxadiene producing *S. cerevisiae* strain, harbouring three copies of taxadiene synthase (*TASY*) genes (Nowrouzi et al., 2020). In a later study, we proposed that the presence of the uncoupling phenomenon potentially fuels the activity of the CYP725A4 enzyme by providing electron and oxygen donors via the radical, reactive intermediates (Nowrouzi et al., 2022). We have also previously shown that increasing the expression of its interacting reductase gene (*POR*) and elevated taxanes concentrations were destructive to the yeast and productivity. This would hinder the use of multiplexing genome editing approaches for increased diterpenoids titre, since the enzymatic-level uncoupling effect would reinforce the oxidative stress (Nowrouzi and Rios-Solis, 2021, Malci et al., 2023). On the other hand, many studies on Taxol® production in *Taxus* spp. displayed a significant impact of biotic and abiotic elicitor application on the terpenoid accumulation (Barrales-Cureño et al., 2022; Escrich et al., 2021; Ketchum et al., 1999; Mirjalili and Linden, 1996). Inspired by this natural phenomenon (Mirjalili and Linden, 1996), we hypothesised that a similar elicitor could potentially stimulate further terpenoid production in yeast. For this reason and to address the associated oxidative stress issue, an adaptive laboratory evolution approach was pursued to adapt the taxadiene-producing yeast cells to increasing concentrations of hydrogen peroxide, with or without dodecane, while growing in galactose-containing complete synthetic mixture to induce the taxanes production in our yeast cells. Using gene expression profiling and exometabolite analysis, we investigated the gene expression profiles of the best-performing evolved colony and the parent strain to examine the alterations in the overall yeast physiology before and during the hydrogen peroxide (H_2_O_2_) induction in galactose-limited chemostats. We also characterised the associated changes in iron and flavin availability, which were found to be critical to optimal P450-reductase for the next enzyme activity in the Taxol® pathway (CYP725A4-POR) (Nowrouzi et al., 2022). The exometabolite results were also integrated into an enzyme-constrained yeast genome-scale model to interpret the flux changes in each condition with more confidence.

## 2. Materials and Methods

### 2-1. Yeast strains

The *S. cerevisiae* strain used for adaptive laboratory evolution was LRS5 (MATa, *leu2-3,112::HIS3MX6-pGAL1-ERG19/pGAL10-ERG8;ura3-52::URA3-pGAL1 MvaSA110G/pGAL10-MvaE* (codon-optimised)*; his3*Δ*1::hphMX4-pGAL1-ERG12/pGAL10-IDI1;trp1-289::TRP1_pGAL1-CrtE(Xanthophyllomyces dendrorhous)/pGAL10-ERG20;YPRCdelta15::NatMX-pGAL1-CrtE*(codon-optimised)*/pGAL10-CrtE::ARS1014a::pGAL1-TASY-GFP::ARS1622::pGAL1-MBP-TASY-ERG20*::ARS1114a::pGAL1-MBP-TASY-ERG20*)*, previously constructed in Nowrouzi et al. study (Nowrouzi et al., 2020), and originated from CEN.PK2-1C (Entian and Kötter, 2007) (EUROSCARF collection). The evolved strain as a result of adaptation to H_2_O_2_ and galactose was named E_LRS5.

### 2-2. Growth media and cultivation platforms

The synthetic defined medium (SD) consisted of 0.79 g/L Complete Supplement Mixture (CSM) powder (Formedium), 5 g/L ammonium sulphate (Fisher Chemical™) and 1.7 g/L yeast nitrogen base without amino acids (Thermo Scientific™) and was supplemented with 2% (w/v) galactose (SDG). The routinely used growth medium was composed of yeast extract (1%(w/v)) (Fisher BioReagents™) and casein peptone (2%(w/v)) (Merck), and was supplemented with 2% (w/v) glucose (YPD) or galactose (YPG) (Thermo Scientific™), for inoculum preparation and production screening, respectively. A 20% (v/v) n-dodecane (ACROS Organics™) was used in all cultivations requiring the taxane quantification to extract the taxanes and prevent their air-stripping.

### 2-3. EC50 determination

A fresh LRS5 glycerol stock was streaked on a YPD agar plate and incubated at 30 °C for two days. A 5-mL YPD culture of one randomly picked colony was prepared. Then, one OD_600_ unit of the overnight was inoculated in 2 mL of YPG medium in a 24-well deep well microplate (Axygen) with a variable range of H_2_O_2_ concentrations to assess the half maximal effective concentration (EC_50_) of this oxidative stress-inducing agent on taxadiene-producing yeast cell. In a parallel run, YPG medium was further supplemented with 20% (v/v) n-dodecane (ACROS Organics™) to address the same question under both oxidative and phase toxicity stresses, representing a more realistic picture of the oxygenated taxanes-producing cells. The biomass accumulations in both conditions were tested at the end of three days. The EC_50_ values were determined using the dose-response fittings calculated by Origin 2021 software.

### 2-4. Adaptive Laboratory Evolution (ALE)

The same LRS5 glycerol stock was again streaked on a YPD agar plate and incubated at 30 °C for two days. Three independent colonies were then each incubated in 5 mL of YPD liquid medium for one day at 30 °C, 250 rpm. The cells were then inoculated at the starting OD_600_= 1 in SDG medium using a six-well microplate (Greiner Bio-One™) and were incubated at 30 °C and 350 rpm in a thermomixer for 48 hours. The Periodic Challenge/Recovery Scheme methodology proposed by Reyes and Kao (Reyes et al., 2014) was adapted to increase the LRS5 tolerance to increasing concentrations of H_2_O_2_ (with or without dodecane) while they were producing taxanes. Briefly, one OD_600_ unit of 48-hour cultures were transferred to a 12-well microplate (Greiner Bio-One™) containing new SDG medium. For each colony, an extra technical replicate was also included. Based on the EC_50_ calculations, the cells were initially challenged with 2.13 mM of H_2_O_2_ (H condition) or 0.46 mM of H_2_O_2_ and 20% (v/v) dodecane (DH condition) for two days and allowed to rest without H_2_O_2_ presence for the subsequent three days, using 50 µL of inoculum for each cultivation round. During the resting period, for DH condition, dodecane was continued to be supplemented in the fresh medium. Parallel extracellular measurement of H_2_O_2_ using Pierce™ Quantitative Peroxide Assay Kit (Thermo Fisher Scientific) ensured that this compound fully entered the yeast. The cells were routinely cross-checked by microscopy for any microbial contamination. The ALE process (two-day challenge and three-day rest) was repeated until no further improvement in the cell growth was observed, when the final H_2_O_2_ resistance concentration reached 10 and 12 mM of H_2_O_2_, with and without dodecane. All colonies were preserved in 50% glycerol at −80 °C for further analysis.

### 2-5. Taxane production assessment of yeast strains

The LRS5 and all evolved strains were incubated in 5 mL of YPD liquid medium at 30 °C, 250 rpm. The 24-hr precultures were then inoculated in 1.5 mL of total YPG medium supplemented with 20% (v/v) dodecane in 48-well FlowerPlates (mp2-labs). The plate was sealed with two layers of gas-permeable membrane (Thermo Scientific™) to avoid dodecane evaporation and product loss. The cells were incubated at 30 °C and 800 rpm for 72 hours. By the end of the cultivation, dodecane was extracted from each triplicate set for further analysis with GC-MS and the biomass accumulations were also measured. The best-performing colony, with the highest taxadiene titre, was labelled E_LRS5 and was used for all the subsequent experiments.

### 2-6. Diterpenoid identification and quantification

A 50 µL of dodecane overlay was extracted from each culture media and analysed by GC–MS using Trace 1300 GC (ThermoFisher Scientific), equipped with TraceGOLD TG-5MS (30 m[×[0.25 mm[×[0.25 μm). The mass spectra in the range of 40–650 m/z was recorded on a Thermo Scientific ISQ Series single quadrupole mass spectrometer at EI ionisation mode. The GC temperature programme began at 120 °C (three minutes) and was then raised to 250 °C at a rate of 20 °C/minute with a three-minute hold time. Xcalibur™ software (ThermoFisher Scientific) was used to process the data. Geranylgeraniol (GGOH, Sigma Aldrich) was used to quantify the GGOH concentration. The diterpene totarol (Cayman Chemical) was used to quantify the taxanes.

### 2-7. BioLector cultivations for growth rate measurement

To determine an appropriate dilution rate for chemostat experiments, BioLector cultivations using a combination of glucose and/or galactose in buffered SD medium (buffered with 0.1 M potassium phthalate monobasic to pH = 6) were performed. The LRS5 and E_LRS5 overnight cultures in YPD were centrifuged at 5000 rpm for 10 minutes, washed with sterile ultrapure water twice, and back diluted to initial OD_600_= 5 in 1 mL total volume of water. A 30 µL of the cell dilution was then used to inoculate a 48 well flower plate (m2p-labs), containing galactose: glucose ratio of 25, 50, 75 and 100 at 1, 2, 3 and 4% total sugar composition. The microplate was sealed with a gas permeable membrane (m2p-labs), and the cultivation was performed in BioLector II microbioreactor at 30 °C and 1000 rpm until the early/full stationary phases were observed for both strains. The maximum specific growth rates for each triplicate were calculated by growthrates R package(Petzoldt, 2020) using a nonparametric smoothing spline function.

### 2-8. Bioreactor cultivations

Fresh glycerol stocks of LRS5 and E_LRS5 strains were streaked on new YPD medium for two days. A single colony of each were then grown in 10 mL of YPD medium at 30 °C, 250 rpm and the whole preculture was transferred to 100 mL of buffered SDG medium (pH = 6) in 500-mL baffled shake flasks for additional 1-2 days until a suitable cell density could be achieved for bioreactor inoculation. The preculture cells were then inoculated at initial OD_600_= 0.1 in triplicates in 1L Biostat Qplus bioreactors (Sartorius Stedim Biotech). The bioreactor cultivations were performed in 500 mL of SDG medium containing 20 g/L galactose (separately autoclaved), 6.7 g/L yeast nitrogen base with ammonium sulphate and without amino acids and 0.79 g/L complete synthetic mixture. The pH was adjusted to six using 2 M NaOH, and two drops of antifoam 204 (Merck) per liter were added to reduce the foaming. The galactose-limited chemostat cultures started once the off-gas CO_2_ started to drop in the batch culture, denoting the depletion of the carbon source. The chemostat cultures were carried out using the same medium supplemented with 10 g/L galactose, at a dilution rate of 0.1 h^-1^, and it lasted for 50 hours (five residence times) for each steady state. Upon RNA and physiological samplings, the steady state II (SSII) started with the same condition, except that the feeding medium also contained H_2_O_2_ at a final concentration of 0.5 mM, added separately after the medium was autoclaved. The bioreactors were run at 30 °C with 600 rpm agitation and airflow of 0.5 SPLM. The pH was controlled to remain at six using 2 M of each NaOH and H₂SO₄. The oxygen uptake rate (*OUR*) and the carbon dioxide evolution rate (*CER*) values were calculated according to the bioreactor manufacturer’s (Sartorius Stedim Biotech) guideline, at 25.4971 L molar volume of gas. The respiratory quotient (*RQ*) was determined by the following formula:

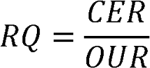

### 2-9. Dry cell weight and extracellular metabolites measurements

All samplings were performed at the same time after removing the dead volume. The dry cell weights and biomass accumulations during each steady state were measured by retrieving 10 mL of culture broth from each reactor. From that, 1 mL was used for OD_600_ measurement and the other 9 mL were filtered using pre-weighed 0.45-µm MCE filter papers (MontaMil ®). This was followed by drying the filter in the microwave for 10 minutes and 2-3 days of desiccation. The extracellular metabolites including ethanol, succinate, acetate, glycerol and pyruvate productions of both steady states and strains were analysed using ion-exchange HPLC. The samples were first filtered using a 0.45 µm filter. The samples were then analysed by injecting them into a Bio-Rad Aminex HPX-87H column at 60 °C. The injection volume was 20 µL and the elution solvent was 5 mM H_2_SO_4_ at a flow rate of 0.6 mL min^-1^. The off-gas ethanol was also included in the total ethanol flux measurement.

### 2-10. Statistical analyses for biomass and flux measurements

All statistical analyses of this study were performed in R version 4.1.3. Two-way analysis of variance (ANOVA) was used to study the effect of the strain type and the continuous culture state (steady states I or II) on carbon recovery and flux changes, using R stats package. The Bonferroni-corrected t-test was performed for the pairwise comparisons, at a significance of adjusted p < 0.05. The biomass and taxanes production by the evolved and parent cells were compared using the Kruskal-Wallis test. For the same analysis, the pairwise comparison against the control condition was performed by Dunnett’s multiple comparisons test at a significance of p < 0.05, using DescTools R package (Signorell et al., 2022).

### 2-11. Sampling for transcriptomics analysis

The RNA-seq samplings were performed by discarding the dead volume followed by retrieving 1-mL culture broth at a minimum of 10 OD_600_ units in Eppendorf tubes, which were pre-cooled at −20 °C. The samples were immediately placed on ice and quickly centrifuged in a pre-chilled centrifuge for two minutes at 19,341 ×g. Upon removing the supernatants, the samples were flash frozen in liquid nitrogen and stored at −80 °C for further analysis.

### 2-12. Total RNA extraction and Library preparation with polyA selection and HiSeq sequencing

Total RNA was extracted using Qiagen RNeasy Plus mini kit following the manufacturer’s instructions (Qiagen). Extracted RNA samples were quantified using Qubit 2.0 Fluorometer (Life Technologies) and RNA integrity was checked using Agilent TapeStation 4200 (Agilent Technologies). RNA sequencing libraries were prepared using the NEBNext Ultra II RNA Library Prep Kit for Illumina following the manufacturer’s instructions (NEB). Briefly, mRNAs were first enriched with Oligo(dT) beads. Enriched mRNAs were fragmented for 15 minutes at 94 °C. First strand and second strand complementary DNAs (cDNAs) were subsequently synthesised. The cDNA fragments were end repaired and adenylated at 3’-ends, and universal adapters were ligated to cDNA fragments, followed by index addition and library enrichment by limited-cycle PCR. The sequencing libraries were validated on the Agilent TapeStation (Agilent Technologies) and quantified using Qubit 2.0 Fluorometer (Invitrogen) as well as by quantitative PCR (KAPA Biosystems) (Figure S3. 1 and Table S3. 1). The sequencing libraries were clustered on a flowcell lane. After clustering, the flowcell was loaded on the Illumina HiSeq instrument (4000 or equivalent) according to the manufacturer’s instructions. The samples were sequenced using a 2×150bp Paired End (PE) configuration. Image analysis and base calling were conducted by the HiSeq Control Software. Raw sequence data (.bcl files) generated from Illumina HiSeq was converted into fastq files and demultiplexed using Illumina’s bcl2fastq 2.17 software. One mismatch was allowed for index sequence identification.

### 2-13. RNA-seq analysis

RNA-seq data was processed using nf-core RNA-seq pipeline version 3.7 (Ewels et al., 2020), run on Nextflow version 20.12.0-edge and singularity configuration profile (Di Tommaso et al., 2017). After pre-processing the reads, they were mapped onto reference Ensembl yeast R64-1-1 genome (available on Amazon Web Services iGenomes) by Salmon (Patro et al., 2017), followed by transcript abundance quantification using the same software with the default parameters. The data quality was checked with multiQC software (Ewels et al., 2016). The tximport R package (Soneson et al., 2015) was then used to aggregate the transcript-level estimates into gene level abundances before differential gene expression using the Wald test implemented in the DESeq2 R package (Love et al., 2014) for both effect-based and pairwise comparisons. The lfcShrink function with ashr shrinkage estimator (Stephens, 2017) was then used to estimate the expression changes more accurately. The differentially expressed genes were filtered by an adjusted p value of < 0.05 and log2 fold change threshold at zero. The pathway enrichment analyses were performed with pathfindR (Ulgen et al., 2019) R package, based on the Kyoto Encyclopedia of Genes and Genomes (KEGG) database (Kanehisa and Goto, 2000) and the p values were adjusted using the default Bonferroni method. The package org.Sc.sgd.db (Carlson, 2019) was used for gene ontology (GO) overrepresentation analysis using clusterProfiler (Yu et al., 2012) package, with p values corrected according to Benjamini and Hochberg’s false discovery rate test (Benjamini and Hochberg, 1995). The gene sets smaller than 10 and larger than 500 genes were ignored for both GO and KEGG enrichment analyses. All RNA-seq analyses were performed on Eddie, The University of Edinburgh’s high-performance computing cluster using 12 cores of 10 Gigabytes RAM memory in Intel® Xeon® Processor E5-2630 v3 (2.4 GHz) processor.

### 2-14. Preparation of models and flux balance analysis

The ecYeastGEM model (GECKO toolbox 2.0 (Domenzain et al., 2022); ecYeastGEM_batch.mat) was used as the yeast model. This enzyme-constrained model already contained 8028 reactions and 4153 metabolites and was manipulated by adding 32 reactions and 24 metabolites with separate forward and reverse, arm and protein exchange reactions for terpenes synthesis and their import and export, to achieve mass balance. For both strains under the stress, the models were extended with hydrogen peroxide intracellular transport and exchange reactions to resemble the stress condition. The coefficients of the enzyme pseudometabolites in each reaction were calculated according to Sánchez et al. (Sánchez et al., 2017), by querying the K_cat_ values from BRENDA database (Chang et al., 2021). When the Kcat values were not directly available for the enzymes, it was calculated using the following formula:

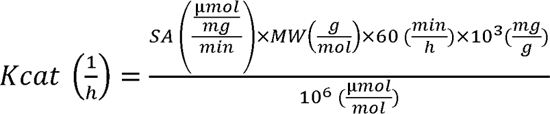

where *sA* is the specific activity and *Mw* is the molecular weight of the enzyme. The heterologous genes in the mevalonate (MVA) pathway were added to their associated reactions in the model. The script for addition of the new reactions and related modifications is available in the figshare repository: https://figshare.com/s/3f00d32c57c2a189d2db; DOI: 10.6084/m9.figshare.20424801.

Four models, two per strain (for each steady state), were then created by constraining the fluxes of off-gas and exo-metabolomes at a fixed growth rate (dilution rate), considering 5% measurement error to avoid model over-constraining. The reactions related to medium components uptakes, terpene export, unmeasured metabolic fluxes and overflow metabolites were set to open with a maximum upper boundary (UB) of 1000 and lower boundary (LB) of - 1000 mmol/gDW/h. The models were then optimised by FBA by setting the non-growth associated maintenance (NGAM) reaction (r_4046) as the objective function. After fixing the NGAM reaction boundaries by the calculated value from the FBA analysis, the flux through cytoplasmic, mitochondrial and peroxisomal thioredoxin reactions (r_0550No1; r_0550No2; r_0551No1; r_0552No1; r_0552No2) were set open to activate the core antioxidant mechanism in all models. The manipulation of the models was finalised by minimising the total enzyme pseudo-metabolite pool exchange flux. All simulations were performed using COBRA Toolbox (Heirendt et al., 2019; Schellenberger et al., 2011) with Gurobi Optimiser (version 9.5.1) as the solver in MATLAB R2021a.

### 2-15. Random sampling of the solution spaces and flux enrichment analysis

The random sampling algorithm implemented in RAVEN toolbox (Agren et al., 2013; Wang et al., 2018) was employed to investigate the flux distributions in each enzyme-constrained model. A total of 10,000 random samples were collected for each model followed by their pairwise comparisons by the Kruskal-Wallis test. Upon normalizing the results by the average CO_2_ production flux value, COBRA Toolbox (Heirendt et al., 2019; Schellenberger et al., 2011) flux enrichment analysis function was used to perform pairwise comparisons of means of random samples from each model to find the changes in fluxes through enrichment by their genes, using hypergeometric 1-sided test and false discovery rate (FDR) correction for multiple testing at a significance level of 0.05 and fold change cut-off = 1. The intersection of the genes enriched by gene expression and flux enrichment in the same regulation direction showed the flux-gene expression correlations. The whole random sampling analysis was performed in MATLAB R2021a on VLX servers on XRDP Linux Remote Desktop, hosted by the School of Engineering at The University of Edinburgh.

## 3. Results & Discussion

### 3-1. Evolving taxadiene-producing yeast on H_2_O_2_ as an elicitor for higher Taxol® precursors production

Earlier we reported that the higher oxygenated taxanes production by *Taxus cuspidata* CYP725A4 and POR was associated with the release of reactive metabolites, potentially acting as oxygen/electron donors while imposing oxidative stress on yeast (Nowrouzi et al., 2022). Therefore, a robust yeast cell factory for producing highly oxygenated taxanes like Taxol® was needed to efficiently address this paradoxical issue. To further examine if the oxidative stress and overall, the toxicity issue, was related to oxygenated-taxanes production, our preliminary experiments tested the effect of increasing concentrations of hydrogen peroxide (H_2_O_2_), a known oxidant agent, on several of our in-house yeast strains. These included the wild-type CEN.PK strain, an engineered CEN.PK strain with overexpressed genes in mevalonate (MVA) pathway (mGTy116) (Reider Apel et al., 2017), taxadiene-producing mGTy116 (LRS5) (Nowrouzi et al., 2020) and oxygenated taxanes-producing LRS5 (LRS6) (Walls et al., 2020) in YPG medium supplemented with dodecane, representing a realistic, normal scenario, under which taxanes are induced and removed *in situ* by an organic overlay. The results revealed the superiority of LRS5 in final biomass compared to mGTy116 and LRS6, where almost half of LRS5 final biomass was retained at 2 mM final H_2_O_2_ concentration (Figure 2A). Hence, we initially concluded that the LRS5 strain might resist the oxidative stress, while mGTy116 showed the weakest resistance, likely due to overaccumulation of toxic prenyl pyrophosphates (Martin et al., 2003). This observation further strengthened our hypothesis that adapting LRS5 to increased oxidative stress would benefit the biomass and taxadiene production over time, as we earlier showed a strong correlation between the biomass accumulation and taxadiene production by LRS5 (Nowrouzi et al., 2020).

**Figure 2.**
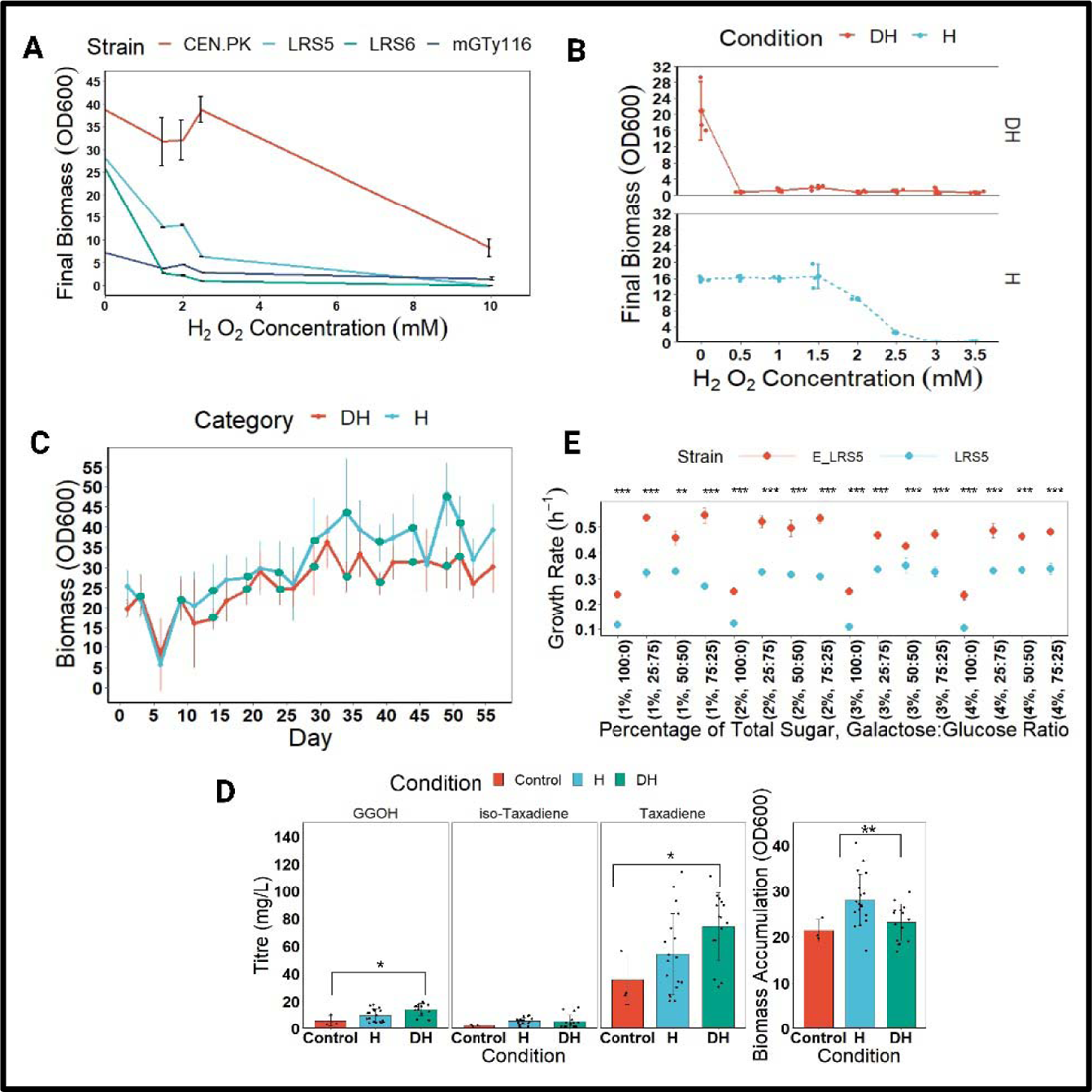
Summary of adaptive laboratory evolution results and small-scale physiological parameters measurements. A) The effect of increased concentration of H_2_O_2_ on the final biomass of selected yeast strains. All strains were cultivated in YPG+20% (v/v) dodecane medium for three days at 30 °C, 450 rpm. CEN.PK: CEN.PK2-1C (Entian and Kötter, 2007); mGTy116: CEN.PK yeast strain with overexpressed genes in mevalonate pathway (Reider Apel et al., 2017); LRS5: mGTy116 harbouring three copies of taxadiene synthase genes (Nowrouzi et al., 2020); LRS6: LRS5 harbouring *Taxus cuspidata* taxadien-5α-hydroxylase (*CYP725A4*), cytochrome P450 reductase (*POR*) and taxa-4(20),11(12)-dien-5α-olO-acetyl transferase (*TAT*) genes for production of oxygenated and acetylated taxanes (Walls et al., 2020). The data show the mean ± SD of biological triplicates. B) EC_50_ determination for LRS5 under dodecane and hydrogen peroxide (DH) or hydrogen peroxide (H) stresses. The LRS5 strain was cultivated for three days at 30 °C, 450 rpm. For all DH conditions, 100% dodecane was utilised. The data show the mean ± SD of biological triplicates. C) Biomass plot for adapting the taxadiene-producing strain (LRS5) to H_2_O_2_ (H) and H_2_O_2_+Dodecane (DH), while growing on a complete synthetic mixture with 2% (w/v) galactose (SDG). The line plots represent the mean ± SD of three biological and two technical replicates. The green dots on each plot show the start of the H_2_O_2_ induction period. The increasing H_2_O_2_ concentrations (concentration increased every 5 days) at H condition were: 2.13, 2.13, 2.63, 3.63, 4.63, 6, 8, 10, 10, 12 and 12 mM. The increasing H_2_O_2_ concentrations at DH condition (increased concentration every 5 days and then fixed) were: 0.46, 0.46, 1, 2, 2, 3, 5, 8, 10, 10 and 10 mM. The values were selected based on the biomass results obtained at the end of each resting period. D) Taxanes and biomass accumulation from parent strain (control: LRS5) and evolved strains (E_LRS5 from H (H_2_O_2_) and DH (H_2_O_2_+Dodecane) conditions) after 72-hour cultivations in YP-galactose medium (2% w/v) at 30 °C. The data show mean ± SD of three biological and two technical replicates for the evolved strains and triplicates of the parent strain (LRS5). The asterisks indicate significant differences by Dunnett’s multiple comparisons test (***: adjusted p < 0.001, **: adjusted p < 0.01, *: adjusted p < 0.05), relative to control conditions (SS II vs. SSI and/or E_LRS5 vs. LRS5). E) Growth rates comparison of the best-performing evolved colony (E_LRS5) and parent strain (LRS5) in SDG medium (pH = 6) containing 1-4% total sugar at 25-100% galactose: glucose ratios. The mean ± SD values of biological triplicates are shown. Figure was created with BioRender.com.

Biphasic culture systems tend to reduce the toxicity and air-stripping of terpenoids produced by the microbial cells (Ajikumar et al., 2010; Nowrouzi et al., 2020; Sarrade-Loucheur et al., 2020). However, the used organic overlay might inevitably be toxic (Jongedijk et al., 2015; Santoyo-Garcia et al., 2022, Galindo-Rodriguez et al., 2023, Santoyo-Garcia et al., 2023). Therefore, to address the oxidative stress with or without the potential overlay phase toxicity under the continuous production of terpenes, two evolution lines were designed to increase the resistance towards the hydrogen peroxide (H) or dodecane and hydrogen peroxide (DH), starting with a suitable dose of H_2_O_2_ in which at least 50% final biomass could be preserved (EC_50_) (Figure 2B). By the end of 56 days, the H_2_O_2_ concentration tolerance was increased by 5.6-fold and 21.9-fold, at H and DH conditions, respectively (Figure 2C).

As illustrated in Figure 2D, the initial screening of the evolved and the parent strain revealed that the taxadiene and its isomer, iso-taxadiene, and geranylgeraniol (GGOH), the dephosphorylated by-product of taxadiene synthase substrate (geranylgeranyl diphosphate; GGPP) (Song et al., 2017), were all increased by 2.3-, 2.5-, and 2.4-fold, and 2.1-,1.5-, and 1.6-fold, in DH and H conditions, respectively, with taxadiene and GGOH production being significantly higher in DH condition compared to the control (p = 0.045 and p = 0.012). The final biomass from both H and DH were found to be significantly different from each other (p = 0.008), but remained relatively similar to that of control. While we achieved superior product titre in DH condition, potentially due to the strains’ pre-adaptation to dodecane or the dodecane acting as an oxygen vector (Blaga et al., 2018), for the sake of simplicity in our study design, the rest of our study only focused on the H condition. Here, we solely further investigated the performance of the best-performing colony evolved from the H condition which we called E_LRS5. The overall taxadiene, iso-taxadiene and GGOH titres by E_LRS5 were 103.3 ± 15.2, 4.00 ± 3.4 and 11.8 ± 2.3 mg/L, which represented a 2.9-(adjusted p = 0.005), 1.7-(adjusted p = 0.001), and 3.3-fold (adjusted p = 0.001) improvement compared to the original LRS5. Similarly, the final biomass accumulation of E_LRS5 also improved by around 1.6-fold to 33.3 ± 3.9 (adjusted p = 0.011), representing a two-fold improvement in the taxadiene yield (3.1 mg/L/OD_600_). All associated chromatograms and mass spectra are provided in Figures S1-3.

To mimic the oxidative stress induction under the controlled conditions, we then sought to study the E_LRS5 and LRS5 physiologies in chemostat cultures, for which we first attempted to determine a dilution rate suitable to both strains. Due to the slow growth of *S. cerevisiae* on galactose compared to glucose (Paulo et al., 2015), the effect of different galactose: glucose composition at 1 to 4% final total sugar concentration was investigated in BioLector microbioreactors using complete synthetic mixture media without any dodecane addition. The results showed that the growth rate significantly increased in all cases in comparison to the LRS5 strain (Figure 2E). Our results were however contrary to Hong et al. study (Hong and Nielsen, 2013), where the maximum specific growth rate of the galactose-adapted yeast strains was lower in a glucose medium. In our study, the maximum growth rate at 3% galactose condition for E_LRS5 (0.25 ± 0.01) exceeded by around 1.6-fold from the maximum specific growth rate of 0.16 h^-1^ reported for CEN.PK 113-7D during the aerobic batch cultivation in a 3% galactose-containing medium (de Jongh et al., 2008). As illustrated in Figures S4-5, notably, the lag phase in the evolved strain using pure galactose was also longer by approximately two-fold with a shorter exponential phase, potentially as a protection mechanism against the stress (Kussell, 2013).

### 3-2. Respiration is an antioxidant defence mechanism in H_2_O_2_ and galactose-evolved taxadiene-producing yeast

Maximum growth rates (µ_max_) of 0.12 ± 0.01 for LRS5 and 0.24 ± 0.01 h^-1^ for E_LRS5 were achieved under 1% galactose addition, therefore our subsequent chemostat experiments were performed at a dilution rate of 0.1 h^-1^. This also presented us with the advantage of eliminating the negative catabolite repression effect of glucose on the induction of galactose promoters, which we previously used to overexpress several of the MVA pathway as well as the taxadiene synthase genes in LRS5 (Nowrouzi et al., 2020), while we appreciate that depending on the strain, the galactose: glucose ratio sensing can play a role in GAL genes induction (Renan et al., 2015). To mimic the stress associated with oxygenated taxanes products in yeast, the LRS5 and E_LRS5 physiologies were studied through RNA-seq, off-gas and exo-metabolites analyses before (steady state I; SSI) and during the oxidative stress induction (steady state II; SSII) in chemostats. To study the impact of evolution and oxidative stress, besides the differential gene expression analysis, enrichment analysis with respect to protein-protein interaction subnetworks (Ulgen et al., 2019) were performed (Tables S2-4; Figures S6. D-E). To this end, at cut-off of log2 fold change (LFC) = 0 and FDR < 0.05, 1607 and 1675 genes were found to be up- and downregulated, respectively, by the evolution (Figure S6. B), from which 90% and 73% were uniquely up- and downregulated, respectively, highlighting the significant impact of adaptive laboratory evolution on the overall superior performance of E_LRS5. This was further supported by the clear genotype separation by principal component analysis (PCA) (Figure S6. A). The number of upregulated and downregulated genes remarkably decreased to 176 and 455, and 223 and 106 for oxidative stress and interaction factors, respectively, denoting the relative robustness of E_LRS5 against oxidative stress.

To investigate the metabolic mechanisms underlying the different physiologies of both strains under the two steady states, an enzyme-constrained yeast genome scale model was used to generate condition-specific models, followed by optimisation towards the maximisation of ATP production (non-growth associated maintenance (NGAM)) and unconstraining the thioredoxin-using reactions, as the heterologous protein production/H_2_O_2_ induction is correlated with increased antioxidant defence mechanism in yeast (Garrido and Grant, 2002). This was followed by random sampling to study the metabolic fluxes with respect to evolution and oxidative stress induction to identify the up- and down-regulated reactions (E_LRS5 vs. LRS5 and their corresponding steady states) at cut-off = 1 and FDR < 0.05, based on the statistically significant overrepresented genes from the flux enrichment analysis (McGarrity et al., 2020).

The carbon recovery profiles revealed that the major contributor to the overflow metabolites was ethanol (SSI-SSII, LRS5: 23.3 ± 1.90 % and 25 ± 0.61 %, E_LRS5: 16.5 ± 1.21 % and 17.4 ± 1.32 %), followed by acetate at a minor level (SSI-SSII, LRS5: 6.64 ± 0.25 % and 7.25± 0.35 %, E_LRS5: 6.72 ± 0.86 % and 7.66 ± 0.58 %) (Figure 3A); During SSI, E_LRS5 produced less ethanol, acetate, CO_2_ and glycerol, in contrast to pyruvate (Table S1). However, this reduction was only significant for ethanol (adjusted p = 0.00016), glycerol (adjusted p = 3.057e-05) and pyruvate (adjusted p = 1.4e-06) production. Succinate production remained negligible for LRS5, while we did not detect this compound in the E_LRS5 medium. The pyruvate dehydrogenase bypass, composed of pyruvate decarboxylase (PDH), aldehyde dehydrogenase (ADHs) and acetyl-CoA synthase (ACSs), plays an intermediate role in the production of the major source of cytoplasmic acetyl-CoA in yeast (Qin et al., 2020) while decarboxylating the pyruvate to ethanol and acetate, also producing NADPH. The higher accumulation of pyruvate in E_LRS5 SSI culture associates well with the downregulation of the genes of pyruvate-catabolising PDC1, PDC5 and DLD1, and the upregulation of pyruvate-synthesising *PYK2* and *AGX1* (McNeil et al., 1994). The lowered acetate flux is probably due to the downregulation of the mitochondrial *ALD5* and cytosolic *ALD6* and their catalysing reactions. Despite this, we observed the downregulation of acetate-using *ACS1* and *ACS2*, where it is likely that the upregulation in *LAT1* and *LPD1* expressions together with others like mitochondrial acetyl-CoA transferase *ACH1* effectively replaced ACSs to produce the acetyl-CoA. At the same time, the absence of succinate in E_LRS5 cultures relates to the general upregulation of succinate dehydrogenases (*SDH1*, *SDH2*, *SDH3* and *SDH4*), which compensated for the slight upregulation of succinate-consuming *LSC2*. While the lower ethanol exchange flux cannot be explained by the upregulation of *ADH3* and *ADH4* and their predicted reaction fluxes and the downregulation of ethanol-metabolising *ADH2*, we can postulate the lowered reaction fluxes from pyruvate consumption mainly be due to decreased flux through pyruvate decarboxylation reactions. The downregulation of glycerol-producing *GPD2* and *GPP1* as well as dihydroxyacetone phosphate-accumulating *GUT2* and *DAK2*, and upregulation of glycerol-consuming *GPD1*, *GPP2* and *GUT1*, justify the reduced glycerol production by E_LRS5.

**Figure 3.**
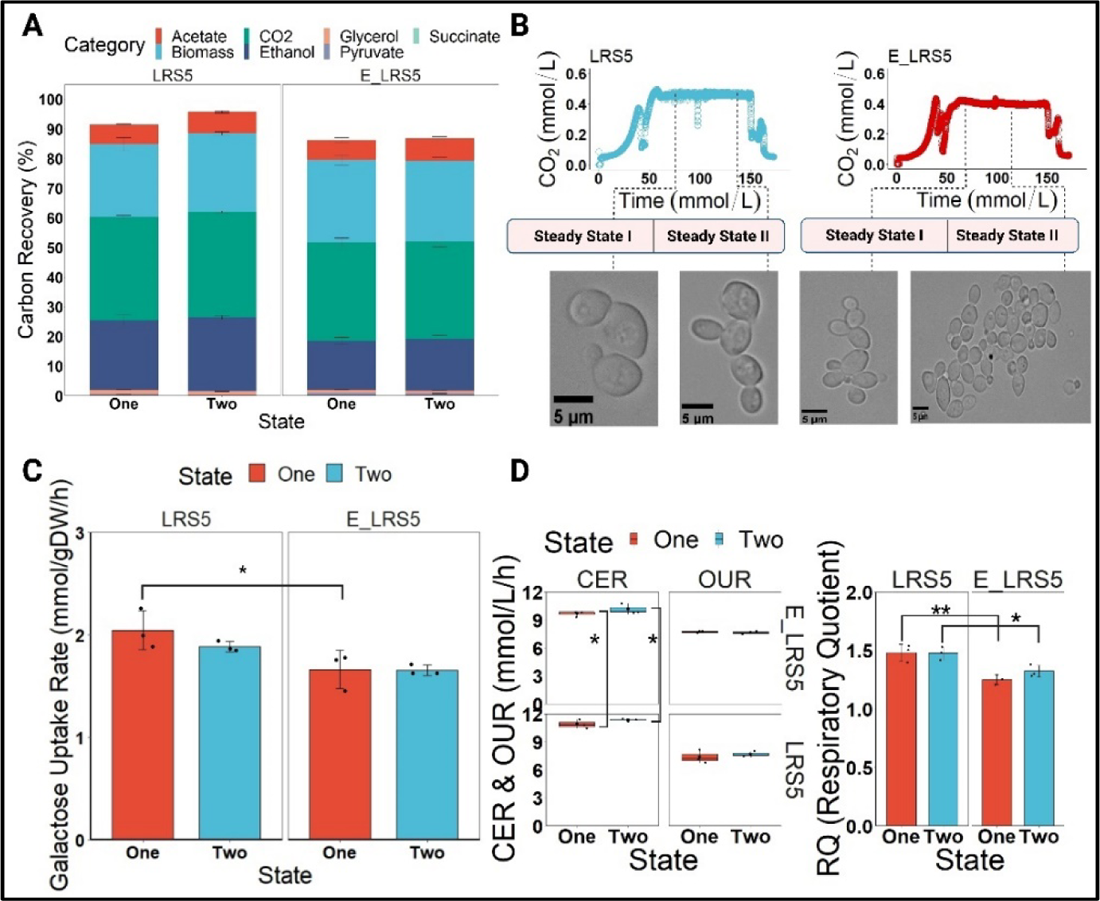
Phenotyping. A) Total carbon recovery for LRS5 and E_LRS5, normalised per mmol of galactose for ethanol, acetate, glycerol, pyruvate, and succinate. The mean ± SD of biological triplicates is shown. The summary of only statistically significant adjusted p values from the t-test (threshold: adjusted p < 0.05) for comparison of product-based carbon recovery is as follows: Glycerol (LRS5: One-E_LRS5: One: 0.0003; E_LRS5: Two-LRS5: Two: 0.009); Ethanol (LRS5: One-E_LRS5: One: 0.001; LRS5: Two-E_LRS5: Two: 0.001); Pyruvate (LRS5: One-E_LRS5: One: 2.22e-07; LRS5: Two-E_LRS5: Two: 6.55e-08). B) Microscopic images of LRS5 and E_LRS5 in continuous cultures before and during oxidative stress induction. The bars represent 5 µm and the images were taken under 100X magnification using a Motic BA310E microscope (Motic, Hong Kong). C) Galactose uptake rates by LRS5 and E_LRS5 chemostat cultures. The cells were cultivated in SDG medium, containing 10 g/L galactose, at a dilution rate of 0.1 h^-1^, during steady state I (SSI), without extra oxidative stress, and with 0.5 mM H_2_O_2_ continuously feeding the cell (steady state II; SSII). The data show mean ± SD of triplicates. D) OUR, CER and RQ plots for LRS5 and E_LRS5. The data show the mean ± SD of biological triplicates. The asterisks indicate significant differences by pairwise t-test (***: adjusted p < 0.001, **: adjusted p < 0.01, *: adjusted p < 0.05), relative to control conditions (SS II vs. SSI and/or E_LRS5 vs. LRS5). Figure was created with BioRender.com.

Upon introduction of the oxidative stress agent, the ethanol and acetate production fluxes further increased, which might be due to the strong downregulation of *ADH2*. It is known that the higher biomass yield can lead to a higher acetate production (Ostergaard et al., 2000), which was in accordance with the higher acetate flux in E_LRS5 upon oxidative induction. Furthermore, the upregulation of *ACS2* and its catalysing reaction flux at E_LRS5 SSII explains the higher accumulation of acetate, similar to LRS5. Compared to LRS5, the E_LRS5 also showed 1.3- and 1.2-fold higher biomass yields per mmol of consumed galactose at SSI (adjusted p = 0.007) and SSII (adjusted p = 0.038), respectively, depicting the better carbon usage in this strain compared to the parent strain (Table S1). An increased TCA capacity can reduce the carbon conversion to ethanol and NADH oxidation into glycerol (Vemuri et al., 2007), explaining the lower glycerol and ethanol accumulation by E_LRS5.

### 3-3. Anticipation and cross-protection mechanisms act to promote the carbon and protein recycling pathways upon oxidative stress re-induction, in H_2_O_2_ and galactose-adapted taxadiene-producing yeast

We noticed a remarkable difference in the yeast morphology between both strains (Figure 3B), where the yeast cells showed more budding scars in the E_LRS5 culture compared to LRS5, and this improved further upon the oxidative stress induction, although similar dry cell weights for E_LRS5 strain were obtained at both steady states (SSI: 2.43 ± 0.15 g/L and SSII: 2.39 ± 0.11 g/L) which might be due to the reduction in cell volume and mitochondrial network expansion (Kim et al., 2020).

Earlier we reported on the filamentous growth of LRS5 due to nutrient starvation (Walls et al., 2022). While we did not observe such elongated filamentous phenotype in our study, one can first relate such morphology to non-quiescence, with characteristics including impaired mitochondrial respiration, decreased trehalose and glycogen conservation and life span extension by chronic oxidative stress (Mohammad et al., 2020). In comparison, however, E_LRS5 depicted a higher respiratory capacity with a 1.2-fold lower RQ value (1.25 ± 0.04 vs. 1.48 ± 0.07; adjusted p = 0.004) (Figure 3D), with similar values at SSII, 1.48 ± 0.06 (LRS5) and 1.32 ± 0.05 (E_LRS5) (adjusted p = 0.039), justifying the lower ethanol accumulation by E_LRS5. We also observed the upregulation of trehalose and glycogen biosynthesis (*GLC3*, *GSY1*, *GPH1*) and downregulation of glycogen hydrolysis, glucoamylases and glucanases like *EXG1*, *BGL2* and β-1,3-glucan synthase *FKS1*. Similarly, *NCW2*, a deletion of which is known to activate the glucan and chitin biosynthesis in the yeast cell wall, was amongst the most strongly downregulated genes (LFC: −2.18) (Queiroz et al., 2020), which interestingly became the most strongly upregulated gene considering the evolution and oxidative stress interaction effect (LFC: 0.69). Similarly, we noticed the general downregulation of the Leloir pathway genes and regulators and reaction fluxes (*GAL1*, *GAL2*, *GAL3*, *GAL7*, *GAL10*, *GAL11* and *GAL80*), which was in accordance with 1.2-fold lower galactose consumption rate in E_LRS5 (1.66 ± 0.2 mmol/gDW/h; adjusted p = 0.038), with almost similar value achieved at SSII (1.65 ± 0.05 mmol/gDW/h) (Figure 3C). The upregulation of *PGM2*, with the enzyme at the glycolysis and pentose phosphate pathway (PPP) branches, was also in line with Hong and Nielsen (Hong and Nielsen, 2013) study, where the adaptive laboratory evolution on galactose resulted in overexpression of *PGM2* towards the trehalose and glycogen biosynthesis. We concurrently observed a strong significant upregulation in maltases like *MAL12*, *MAL32*, isomaltase *IMA1*, glucose-producing glycogen phosphorylase *GPH1* and most hexose transporters (*HXT17*, *HXT6*, *HXT13* and *HXT7*). Considering both evolution and oxidative stress, however, the *HXT6* gene was downregulated, without any other changes in the other stated genes. Hence, we can postulate that to combat the carbon starvation at the next oxidative stress induction, E_LRS5 preferentially invested in recycling the intracellular carbon instead of taking up more galactose and for this, it tended to synthesise the long-chain carbons as protecting and carbon-storing agents (Kocaefe-Özşen et al., 2022). Further to this hypothesis, except the upregulation of *TDH2*, *PGK1*, *PGM2*, *GPM1*, *GPM3* and *PYK2*, we observed the downregulation of glycolysis pathway genes, and the reaction fluxes, as well as the upregulation of non-oxidative PPP and its fluxes due to downregulation of *PGI1* (Qin et al., 2020; Ralser et al., 2007). Indeed, lower glycolysis has been associated with increased oxidative phosphorylation to promote the respiration (Dai et al., 2018) and to improve the stress tolerance (Martínez et al., 2016). This well matches the significant upregulation of the majority of oxidative phosphorylation complexes by evolution, as opposed to both evolution and oxidative stress induction, which only slightly upregulated a few corresponding genes including *ATP14*, *COX8* and *VMA6*, while downregulated *PMA1* and *ATP2*, explaining the slightly higher RQ value of E_LRS5. Clearly, upon introducing the exogenous H_2_O_2_, the correlated flux through glycolysis and the oxidative PPP remained metabolically unchanged, highlighting the robustness of the E_LRS5 upon the oxidative stress re-exposure.

Secondly, we noticed the strong upregulation of the nutrient starvation and low-pH induced genes (Zuzuarregui and del Olmo, 2004) *SPI1* and *YGP1* (LFC: 1.44 and 2.06, respectively). SPI1 is a weak acid stress-induced enzyme which through the cell wall remodelling, it improves the cell viability and minimises the plasma membrane damage (Simões et al., 2006) and its overexpression leads to pseudohyphal growth (Cardona et al., 2012). The overexpression of *YGP1* which is also a yeast cell wall glycoprotein has been linked to a fluffy, non-smooth yeast colony morphology that more readily forms filaments (Kuthan et al., 2003).

Thirdly, it is also likely that the signalling pathways have also played a part as according to Hong and Nielsen (Hong and Nielsen, 2013), the evolution on galactose caused mutations in RAS-Protein Kinase A (Ras-PKA) signalling pathway, and not in the galactose pathway. Likewise, we observed the strong upregulation of its flocculin-encoding gene *FLO11* by evolution, while several Mitogenl7activated protein kinase (MAPK) pathway genes like *SLT2*, *SSK1*, *HOG1*, *YPS1*, *SWI6*, *CDC42*, *MID2*, *STE11*, *ATG33*, and *ROM2* were downregulated opposed to the effect of evolution and oxidative stress.

Fourthly, the sensing of nutrient limitation has potentially activated the MAPK cascade sub-pathway known as the cell wall integrity (CWI) pathway, which regulates the filamentous growth pathway (Birkaya et al., 2009). In this regard, however, several cell surface sensors (*SLG1*, *WSC3* and *MID2*) were downregulated, and *PTC1*, with a role in the CWI reduction (Bi and Park, 2012; Sanz et al., 2022), was upregulated. Similarly, *PRM5*, known as a marker for CWI activation involved in cell wall biogenesis induction, was one of the most strongly downregulated genes due to evolution (LFC= −2.63), decreasing the possibility of CWI being involved in E_LRS5 filamentous growth. On the other hand, the secretory pathway stress and damage to ER quality control can also trigger the CWI pathway and activate the unfolded protein response (UPR) to reduce the cell wall damage and increase the secretory pathway folding capacity (Jiménez-Gutiérrez et al., 2020). This potentially explains the moderate upregulation of genes related to protein sorting into endosome and vacuolar, Golgi and exocyst trafficking as well as the misfolded-protein targeting heat shock proteins HSP42 and HSP26 and oxidative-stress defeating HSP70 isoform, *SSA3* (Gupta et al., 2018; Specht et al., 2011). Indeed, the protein folding and secretion are improved by UPR and heat shock response (HSR), whereby the HSR activates the ubiquitin-proteasome and chaperone to relieve the cell stress (Jiménez-Gutiérrez et al., 2020).

Opposed to the sole evolution effect, the cumulative effect of both evolution and oxidative stress induction caused the upregulation of the endoplasmic reticulum protein processing pathway, ubiquitin-mediated proteolysis and endocytosis pathway genes. Hence, apparently, the evolved yeast has devised different protection strategies to deal with the misfolded proteins. It would then be intriguing to imagine that the exogenous oxidative stress application resulted in increased protein degradation to compensate for this extra burden on E_LRS5 which could also contribute to more ROS formation. This clearly shows a combination of the cross-protection and anticipation likely occurred to improve the microbial fitness against the stressor, potentially in favour of the diterpenoids formation. In this regard, compared to LRS5, E_LRSII showed a 1.1-fold higher NGAM flux at SSI, showing that this strain spends higher energy producing the correctly folded protein and potentially reducing the misfolded protein fraction. On the other hand, increased respiration brought forward more ATP yield and hence a balance between ATP production and ribosome biosynthesis was needed. In this regard, we noticed that the cytosolic ribosome biogenesis genes were downregulated, while the mitochondrial ribosomal biogenesis genes were upregulated by evolution, the opposite of which was observed by the cumulative effect of both evolution and oxidative stress induction, likely due to the already high availability of them due to evolution. Indeed, respiration is less proteome-efficient than fermentation (Elsemman et al., 2022) and ribosome synthesis can limit the growth (Pfeiffer and Morley, 2014). It is however worth noting that the yeast produces a surplus of ribosomes during stressful conditions to quickly use them for fast growth upon shifting to the normal condition (Metzl-Raz et al., 2017). Despite not having studied the proteomes of the studied conditions, apparently, the ribosome biogenesis likely favoured the respiratory processes rather than the metabolic pathways in E_LRS5, an approach LRS5 strain followed to respond to three different factors of taxadiene, galactose and oxidative stress while minimising its enzymatic costs to further couple its productivity with its growth. We can therefore postulate that the enzymatic contents related to taxadiene synthesis were favoured with sufficient ribosome biosynthesis, lipid and vesicular machinery while reducing the proteome allocation costs for metabolic pathways like glycolysis, resulting in E_LRS5 faster growth rate at our initial experiments (Dai et al., 2018).

### 3-4. The decreased purine biosynthesis and increased cell cycle arrest and hydrophilins expressions likely promoted the antioxidant defence mechanism and longevity in H_2_O_2_ and galactose-adapted yeast

We also noticed the downregulation of gene expression and reaction flux through glycine-cleaving GCVs in the folate cycle, which denote the nitrogen source sufficiency in E_LRS5 SSI, preventing the glycine catabolism (Piper et al., 2002). Likewise, there was a drop in purine biosynthesis, likely to prevent the excess ATP production, and this was particularly more evident for E_LRS5 SSII than that of LRS5. Previous studies have illustrated that purine starvation results in increased oxidative stress tolerance and the cell cycle arrest (Kokina et al., 2019; Ozolina et al., 2017). We can then hypothesise that the downregulated purine biosynthesis together with any nutrient starvation and galactose supplementation, all have contributed to the improved survivability and potential cell cycle arrest to promote yeast longevity and the antioxidant events (Petti et al., 2011), representing an active budding phenotype and causing the activation of the non-respiratory mitochondrial genes (Petti et al., 2011). As a result, mitophagy, a quality control process for the degradation of damaged mitochondria to suppress the mitochondrial ROS accumulation (Kanki et al., 2015), was significantly enriched considering both evolution and oxidative stress effects. Moreover, the evolution upregulated the glutaredoxins and glutathione S-transferases, as the yeast’s major antioxidant machinery, together with thioredoxins and glutathione/ glutaredoxins (Grant, 2001). In this regard, the mitochondrial thioredoxin reductase *TRX3*, was significantly upregulated, indicating the long-term adaptation mainly affected the mitochondrial thioredoxin system (Cheng et al., 2019), while both evolution and oxidative stress likely affected the cytosolic thioredoxin system by upregulating *TRX1*. Similarly, the sodium and potassium ATPases, known to protect against the induced cell death (Hoeberichts et al., 2010) were amongst the most highly downregulated genes by the evolution (LFC: −1.67, −1.91, −1.85 and −1.12, respectively). Likewise, hydrophilin genes (induced under the dehydration stress) (*SIP18*; LFC: 3.17, *STF2*; LFC: 0.54, *GRE1*; LFC: 1.91) were strongly upregulated by evolution. *SIP18* is known to reduce the H_2_O_2_-induced yeast apoptosis (Rodríguez-Porrata et al., 2012) and *GRE1* and *STF2* can confer antioxidant capacity as they have been associated with increased cell viability upon restoring the rehydration (López-Martínez et al., 2012).

### 3-5. Taxadiene has antioxidant properties and the improved TCA cycle activity likely provides the required NADPH for diterpenoids production by H_2_O_2_ and galactose-adapted yeast

Overexpressing the phosphoketolase and transaldolase in PPP downstream has been shown to increase the NADPH capability (R. Chen et al., 2022). Also, acute oxidative stress induction was shown to inhibit glycolysis, but increasing the PPP metabolism in Hayes et al. study (Hayes et al., 2020). NADPH is also a redox sink, whereby its consumption elevates during oxidative stress as a means of detoxification (Pollak et al., 2007) and the Taxol® cytochrome P450 enzymes like CYP725A4 also rely on its electrons to activate the dioxygen molecule for diterpenoids production. Hence, we could assume that the oxidative stress likely benefits the Taxol® P450s activity in future aside from the other advantages we stated earlier.

As illustrated in Figure 4, herein, we noticed the downregulation of NADPH-producing *ZWF1*, *ALD6*, *ALD5*, *GND1*, *TKL1* and *RPE1*, the deletion of the last three been reported to increase the sensitivity to hydrogen peroxide (Thorpe et al., 2004). In contrast, the other NADPH-producing enzymes, cytosolic IDP2 and peroxisomal IDP3 isocitrate dehydrogenases belonging to the TCA cycle were upregulated, denoting that the TCA cycle is responsible for the NADPH generation in the evolved strain, where most TCA cycle genes and reaction fluxes were upregulated. Also, the evolution caused a general downregulation of arginine, alanine, lysine, leucine, and tryptophan biosynthesis genes. It is known that proline, serine and alanine catabolism and decreased arginine biosynthesis can help in ATP regeneration through TCA cycle (Liang et al., 2014) and proline is known to act as the oxidative and osmotic stress protectant (Liang et al., 2014). Also, decreased arginine and sulphur amino acid biosynthesis are known to promote lipid biosynthesis and oxidative stress reduction for recombinant protein production (X. Chen et al., 2022). The polyamines exporter protein-encoding *TPO1* and ornithine decarboxylase *SPE1* were also downregulated, both known to be activated during oxidative stress to increase the lysine uptake from medium to free up NADPH for antioxidant system (Olin-Sandoval et al., 2019). This denotes the sufficiency of NADPH cofactor in E_LRS5 strain as indicated previously. However, we noticed an upregulation of the sulphur assimilation pathway genes acting upstream to the antioxidant enzymes. Therefore, our previous assumption for increasing the oxidative stress as a proxy for higher NADPH availability was partly true, as the upregulation of the NADPH-requiring amino acid biosynthesis and sulphur assimilation pathways both pinpoint to the NADPH inadequacy (Ask et al., 2013).

**Figure 4.**
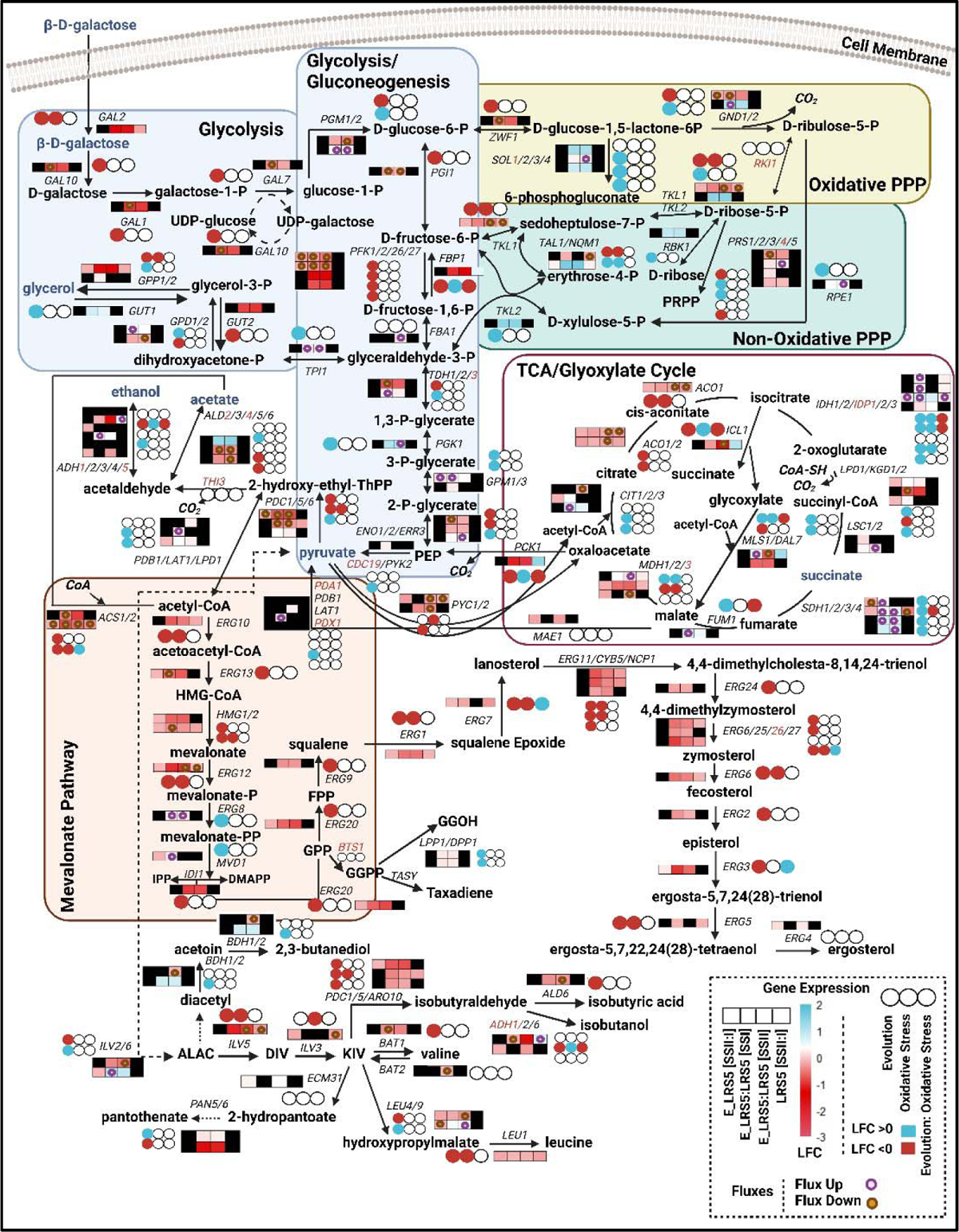
Effect of the adaptive laboratory evolution and oxidative stress on the central carbon metabolism gene expression and flux changes. The mevalonate pathway produces the precursors to terpene and ergosterol biosynthesis. Significant changes to the gene expression and fluxes (mean of n = 10,000 random samples) were selected at log2 fold change = 0 and fold change = 1, respectively, with FDR < 0.05. The heatmap boxes show the following pairwise gene expression and flux comparisons from left to right (changes with regards to oxidative stress induction in E_LRS5; changes with regards to evolution only (during steady state I); changes with regards to evolution effect during steady state II; changes with regards to oxidative stress induction in LRS5). The black colour in the heatmaps shows no significant changes to the expression of the genes at the specific contrast level. The differential gene expressions were also examined by the effect of evolution, oxidative stress and their interplay and are displayed by triple circles in the order of left to right, where the upregulation and downregulation of the genes are shown with blue and red colours, respectively. Genes without significant expression change at any comparison were highlighted in red. The white colour in the triple circles denotes no significant changes to the expression of the genes with respect to condition variables. Where the flux changes were increased or decreased compared to the control level by their genes, it was shown with purple and yellow circles, respectively. The -P stands for phosphate for all metabolites. Figure was created with BioRender.com.

As the expression of their genes remained unchanged due to both evolution and oxidative stress effects, it is therefore likely that the NADPH sources would still be abundant for CYP725A4 activity, however, further experiments would still be needed. We, nevertheless, found a significant general downregulation in the early MVA pathway, forming the backbone of the Taxol® precursors, and NADPH-requiring fatty acid biosynthesis genes expression and reaction fluxes, except for *ERG8* and *MVD1*, catalysing the conversion of mevalonate-P mevalonate-PP and to IPP/DMAPP. Nevertheless, both evolution and oxidative stress induction upregulated the expression of *ERG3*, *ERG7* and *ERG27* (Figure 4). Besides giving strength and fluidity to yeast cell membranes, ergosterols have been reported to increase yeast resistance towards different lignocellulosic inhibitors (Endo et al., 2008). The downregulation in this pathway was contrary to our previous strategy of overexpressing its genes for improving the taxadiene titre (Nowrouzi et al., 2020) as well as to Hong et al. study (Hong et al., 2011), where galactose adaptation upregulated the ergosterol pathway. Furthermore, as the excess NADPH is known to be spent in the ergosterol pathway to counteract its harmful effect, there is a possibility that either E_LRS5 does not over-accumulate the NADPH, or the evolved cell prioritises its use for other biological processes. In this regard, we however observed that *LPP1* and *DPP1*, catalysing the conversion of GGPP to GGOH (Song et al., 2017), were upregulated by the evolution, possibly indicating the GGPP overaccumulation (Nowrouzi et al., 2020) despite the MVA pathway downregulation. Since we observed higher GGOH and taxadiene titre in E_LRS5, it is plausible that yeast already possessed a more active MVA pathway which necessitated its strong downregulation to reduce the toxicity of its products. A recent study by Jeong et al. (Jeong et al., 2021) also mentioned GGPP toxicity in *E. coli; therefore*, it is likely that the increased GGOH titre was to counteract the excess GGPP toxicity.

### 3-6. The oxidative stress re-induction in H_2_O_2_ and galactose-adapted taxadiene-producing yeast likely promotes the flavin formation in favour of CYP725A4

Taxadiene is the substrate to Taxol® CYP725A4 and its interacting reductase POR, each functioning through their prosthetic groups, being heme and flavins, respectively. In addition, the efficient assembly of the cofactor-requiring components of the electron transfer chain (ETC), active in mitochondrial respiration, relies on both mitochondrial flavin adenine dinucleotide (FAD) import, and heme synthesis. We, therefore, studied the changes to biosynthetic pathways of these cofactors to answer if E_LRS5 suited the integration of the later P450 enzyme(s).

The flavin-dependent enzymes, mainly located in mitochondria, constitute about 1.1% of the total yeast proteome, with the majority requiring FAD and the rest depending on flavin mononucleotide (FMN), both formed initially from riboflavin (Gudipati et al., 2014). Briefly, the riboflavin biosynthesis forms through the conversion of guanosine triphosphate (GTP) or ribulose 5-phosphate to its intermediate or through direct riboflavin uptake from the medium via plasma membrane flavin transporter MCH5 (Spitzner et al., 2008). This is followed by the enzymatic steps encoded by *RIBs*, and then the activity of the riboflavin kinase (FMN1) to synthesise the FMN and its conversion into FAD in mitochondria by FAD synthase (FAD1) and its export into the extramitochondrial phase via FLX1 FAD transporter enzyme (Bafunno et al., 2004).

The Flavin cofactor availability is of extreme importance, where for instance the increased phenolic acids titre was achieved by overexpression of *MCH5*, *RIB1* and *FLX1* (R. Chen et al., 2022). While riboflavin has been reported to increase the resistance against the H_2_O_2_ (Maslanka and Zadrag-Tecza, 2022), in our study, except *RIB7* and its reaction flux that was upregulated, other RIBs including *RIB1*, *RIB4* and *RIB5* as well as the putative endoplasmic reticulum-associated FAD transporters *FLC1* and *FLC2* and the mitochondrial FAD transporter *FLX1*, were all downregulated by evolution, potentially denoting the lower availability of the mitochondrial FAD in E_LRS5 compared to LRS5. In contrast, the interaction of the evolution and oxidative stress both resulted in increased expression of *RIB4* and *RIB5*, potentially highlighting the overaccumulation of FMN and FAD precursors by the evolution that would suffice at the SSII (Figure 5A). Also, at SSII, only *RIB5* and *FMN1* were upregulated, potentially to counteract the extra stress burden that needed to be compensated with the FMN availability (Chen et al., 2020). While our earlier study did not show any significant effect from FMN on diterpenoid production, others have reported on its remarkable toxicity reduction capability in yeast (Chen et al., 2020; X. Chen et al., 2022). Hence, we can speculate that E_LRS5 likely better serves the expression of Taxol® P450 genes like CYP725A4, despite observing no changes in *FAD1* expression and its promoting effect on increased biomass and diterpenoids production and electron transfer (Gudipati et al., 2014). However, there is also a possibility that the FAD was already highly available, and the earlier steps were rate-limiting for its higher production.

**Figure 5.**
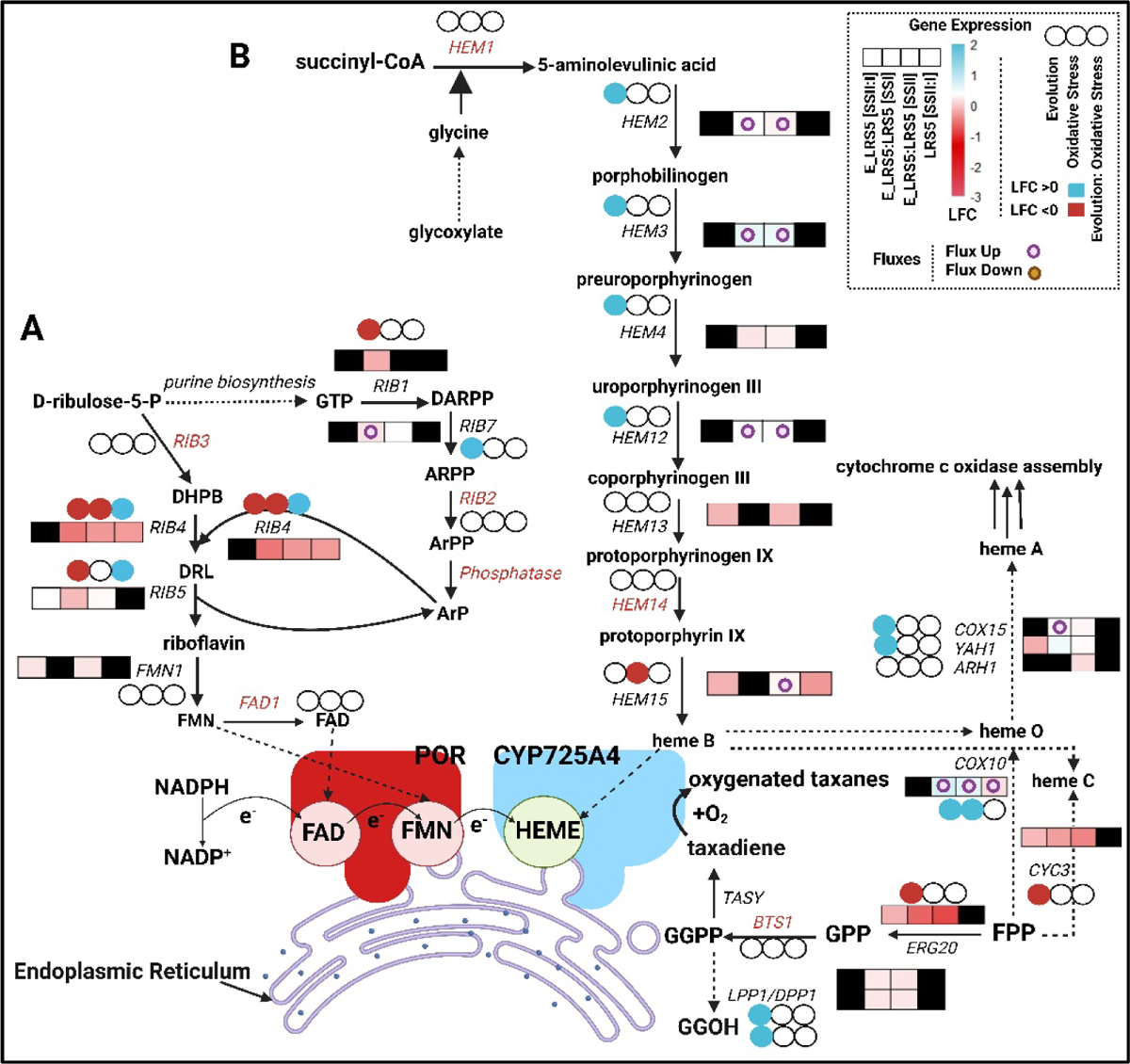
Riboflavin and heme synthesis flux and expression profile alterations with respect to evolution and oxidative stress induction for A) flavin and B) heme biosynthesis. Significant changes to the gene expression and fluxes (mean of n=10,000 random samples) were selected at log2 fold change = 0 and fold change = 1, respectively, with FDR < 0.05. The heatmap boxes show the following pairwise gene expression and flux comparisons from left to right (changes with regards to oxidative stress induction in E_LRS5; changes with regards to evolution only (during steady state I); changes with regards to evolution effect during steady state II; changes with regards to oxidative stress induction in LRS5). The black colour in the heatmaps show no significant changes to the expression of the genes at the specific contrast level. The differential gene expressions were also examined by the effect of evolution, oxidative stress and their interplay and are displayed by triple circles in the order of left to right, where the upregulation and downregulation of the genes are shown with blue and red colours, respectively. Genes without significant expression changes at any comparison were highlighted in red. The white colour in the triple circles denote no significant changes to the expression of the genes with respect to condition variables. Where the flux changes were increased or decreased compared to the control level by their genes, it was shown with purple and yellow circles, respectively. The -P stands for phosphate and e^-^ stands for electron.

### 3-7. The oxidative stress re-induction in H_2_O_2_ and galactose-adapted taxadiene-producing yeast decreased the iron overaccumulation and heme b degradation

In *S. cerevisiae*, the heme synthesis starts with succinyl-CoA and glycine condensation or through importing iron into mitochondria under the iron deficiency (Li et al., 2020). We already observed an upregulation in the TCA cycle succinyl-CoA-producing *LPD1* and *KGD2*, as well as the AGX1, catalysing the formation of glycine from glyoxylate.

We set out to investigate the expression changes of genes directly involved in heme b synthesis for its incorporation into the CYP725A4 enzyme. The heme b synthesis goes through the porphyrin metabolism pathway, which was also found to be significantly enriched by the evolution, where the expression and reaction fluxes of *HEM2*, *HEM3*, *HEM4* and *HEM12*, with enzymes involved in synthesising the intermediates to heme b synthesis were all upregulated (Figure 5B). Similar to the metabolic shift from fermentation to respiration that induces the HAP transcription factor to stimulate the heme-requiring respiratory proteins and ETC (Zhang et al., 2017), *HEM2,3* and *12* genes are known to activate the induction of the TCA cycle and oxidative phosphorylation to source more succinyl-CoA for heme synthesis (Zhang et al., 2017), which aligns well with our previous observation: more active respiration in E_LRS5, compared to LRS5. Similarly, heme b-consuming enzymes encoded by heme A: farnesyltransferase *COX10*, heme a synthase *COX15* and mitochondrial ferredoxin *YAH1* were all upregulated by evolution. However, upon introduction of H_2_O_2_ in E_LRS5 culture, no expression change was detected, except that the heme b-synthesising *HEM1*, *HEM13*, *HEM15* and heme b-consuming *YAH1* expressions were decreased. Indeed, while heme synthesis can benefit the CYP725A4 expression in the Taxol® pathway, we earlier showed that the heme, be it ready heme or heme precursor like 5-aminolevulinate, can negatively result in decreased biomass and diterpenoid formation (Nowrouzi et al., 2022). This, therefore, justifies the increased reaction flux and expression of *HEM2* to consume these iron sources. Interestingly, comparing the LRS5 and E_LRS5 gene expressions upon oxidative stress induction, no changes to *COX10* expression occurred in E_LRS5 and apparently, minimising the heme b degradation can render E_LRS5 a possibly better cell factory for expressing the Taxol® P450 enzymes. Here, we also observed the upregulation of *HAP2*, *HAP3*, *HAP4* and *HAP5* genes due to evolution. *HAP1*, known to induce the expression of the ergosterol biosynthesis and superoxide dismutase genes, was however downregulated, potentially explaining the downregulation of the ergosterol pathway. We, however, can speculate that despite the downregulation of *HAP1*, the upregulation of *HAP2-5* regulated the flux TCA cycle and ETC enzymes (Zhang et al., 2017), while being in accordance with more respiratory status at E_LRS5 during steady state I (SSI). From the other perspective, we can argue that the iron economisation was towards prioritising the expression of H_2_O_2_-degrading iron-containing catalases like heme b-containing *CTT1* (Cassanova et al., 2005). Similarly, *CCC1*, known to be activated to store the excess iron and defeat the iron toxicity (Li et al., 2017, 2001; Li and Ward, 2018) under the excess cytosolic iron availability, was downregulated by evolution, pinpointing to potential iron limitation in the cytosol. Despite this, considering the effect of both evolution and oxidative agent, we observed its upregulation, likely to minimise the ROS generation by excess iron.

## 4. Conclusions

Earlier studies revealed the presence of oxidative stress in *S. cerevisiae* cell factories expressing *Taxus* sp. *CYP725A4* and its cognate reductase gene *POR*, hampering the production of the highly oxygenated anticancer drug, Taxol® precursors in yeast. Using adaptive laboratory evolution, we evolved a taxadiene-producing yeast strain with increased tolerance against hydrogen peroxide (H_2_O_2_) while growing on a galactose-containing medium. The best-performing evolved strain showed three-fold higher taxadiene titre, and later, decreased overflow metabolites production in galactose-limited chemostat cultures. To mimic the oxidative stress induction, the exometabolic and expression profiles of the best-performing evolved strain and the parent strain before and during the H_2_O_2_ induction were studied. Integrating these data with an enzyme-constrained metabolic model showed that the evolution caused the strong upregulation of the mitochondrial respiration as well as the downregulation of the glycolysis and mevalonate pathway, the latter forming the geranylgeranyl diphosphate (GGPP) as the taxadiene synthase substrate. This likely indicated that the antioxidant properties of taxadiene caused its higher production or this was to defeat the GGPP toxicity as reported earlier, all of which potentially resulted in high yeast budding and cell cycle arrest as a defence mechanism. The higher respiration was associated with the elevated upregulation of mitochondrial ribosomal and downregulation of the cytosolic ribosomal genes. The upregulation of the TCA cycle and antioxidant machinery and the downregulation of amino acid production, likely pinpoint to the higher NADPH in the evolved strain, which also acts as the cofactor to cytochrome P450 enzymes of the Taxol® pathway. The oxidative stress induction also downregulated the heme b biosynthesis and upregulated the flavin biosynthesis, which were earlier reported to improve Taxol® early diterpenoids production. On the other hand, the oxidative stress likely caused higher protein misfolding in the evolved strain, while the re-introduction of the oxidative stress induced the proteasomal degradation, as was with storing the branched oxidant-protecting molecules like trehalose and glycogen and hydrolysing them upon H_2_O_2_ re-induction. This points to the existence of both cross-protection and anticipation mechanisms that contributed to greater microbial fitness, being beneficial to Taxol® cytochrome P450 enzymes in the long term. Clearly, the presence of mutation(s) have contributed to the better fitness of the evolved strain, which can be further investigated with genome sequencing and reverse engineering in future studies following systematic approaches (Malci et al. 2022b), nevertheless, these were out of the scope of this study. As of now, the expression of Taxol® pathway genes are conveniently controlled by galactose induction in yeast. Hence, changes to cell factory design are suggested to only activate the *CYP725A4*-*POR* expression upon achieving sufficient intracellular taxadiene level and to allow for yeast to optimise its resources timely and stepwise. On this matter, there is still a need to obtain purified taxadiene to first enable the faster development of specific taxadiene-responsive promoter-based biosensors.

## 5. Data, Materials, and Software Availability

Upon obtaining formal written consent from the first author, all evolved strains generated in this study can only be made available for non-commercial purposes (Nowrouzi, 2022). The log2 fold changes and adjusted p values for the genes discussed/illustrated in this manuscript are available at https://figshare.com/s/3717f4b900bc19124ea7 (RNA-seq) and https://figshare.com/s/7bd488ddf6b4516876d8 (flux enrichment analysis). The gene expression datasets are available at Gene Expression Omnibus. All other generated datasets will be made available on the supplementary information or upon request from the corresponding author.

## Acknowledgements and Funding

The authors would like to thank Mark Lauchlan and Stuart Martin at The School of Engineering, University of Edinburgh, UK for their technical support with GC-MS analysis. Thanks to Tina Johansen at the DTU Fermentation Core, DTU, Denmark for her assistance with HPLC analysis. This work was supported by The University of Edinburgh’s Principal’s Career Development Scholarship, Federation of European Microbiological Societies Research and Training grant (FEMS-GO-2021-044), European Molecular Biology Organization Scientific Exchange Grant (9232), British Council - Metabolic Engineering grant (Grant number: R46889), Engineering and Physical Sciences Research Council (Grant number EP/R513209/1) and the Novo Nordisk Fonden, within the framework of the Fermentation Based Biomanufacturing Initiative (Grant number NNF17SA0031362) and the AIM-Bio project (Grant number NNF19SA0057794).

