## supplementary information for "Rewiring *Saccharomyces cerevisiae* metabolism for optimised Taxol® precursors production"

### Supplementary File

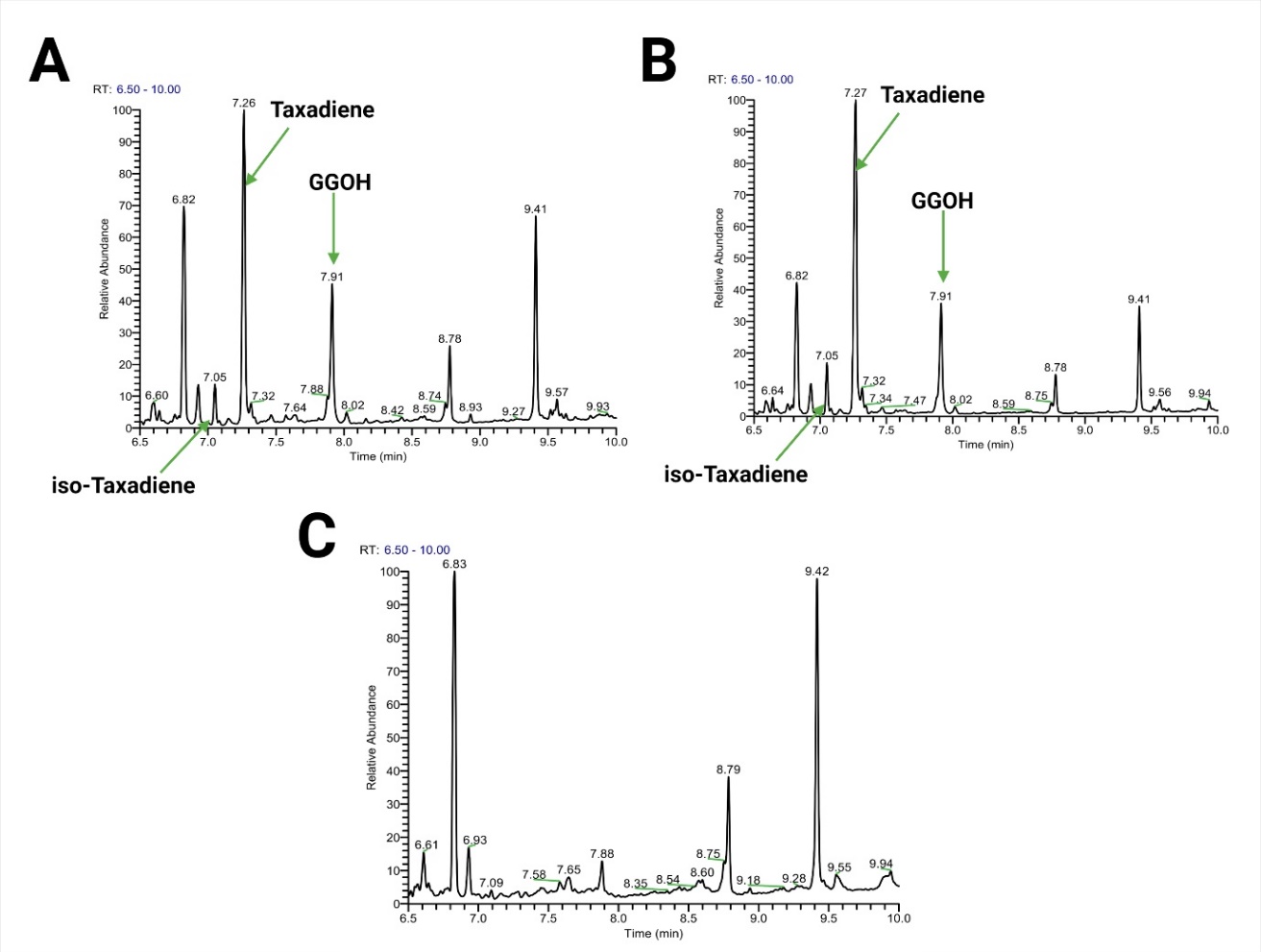

Figure S1. **Sample chromatograms.** A) LRS5; B) E_LRS5; C) Blank (pure dodecane). Figure was created with BioRender.com.

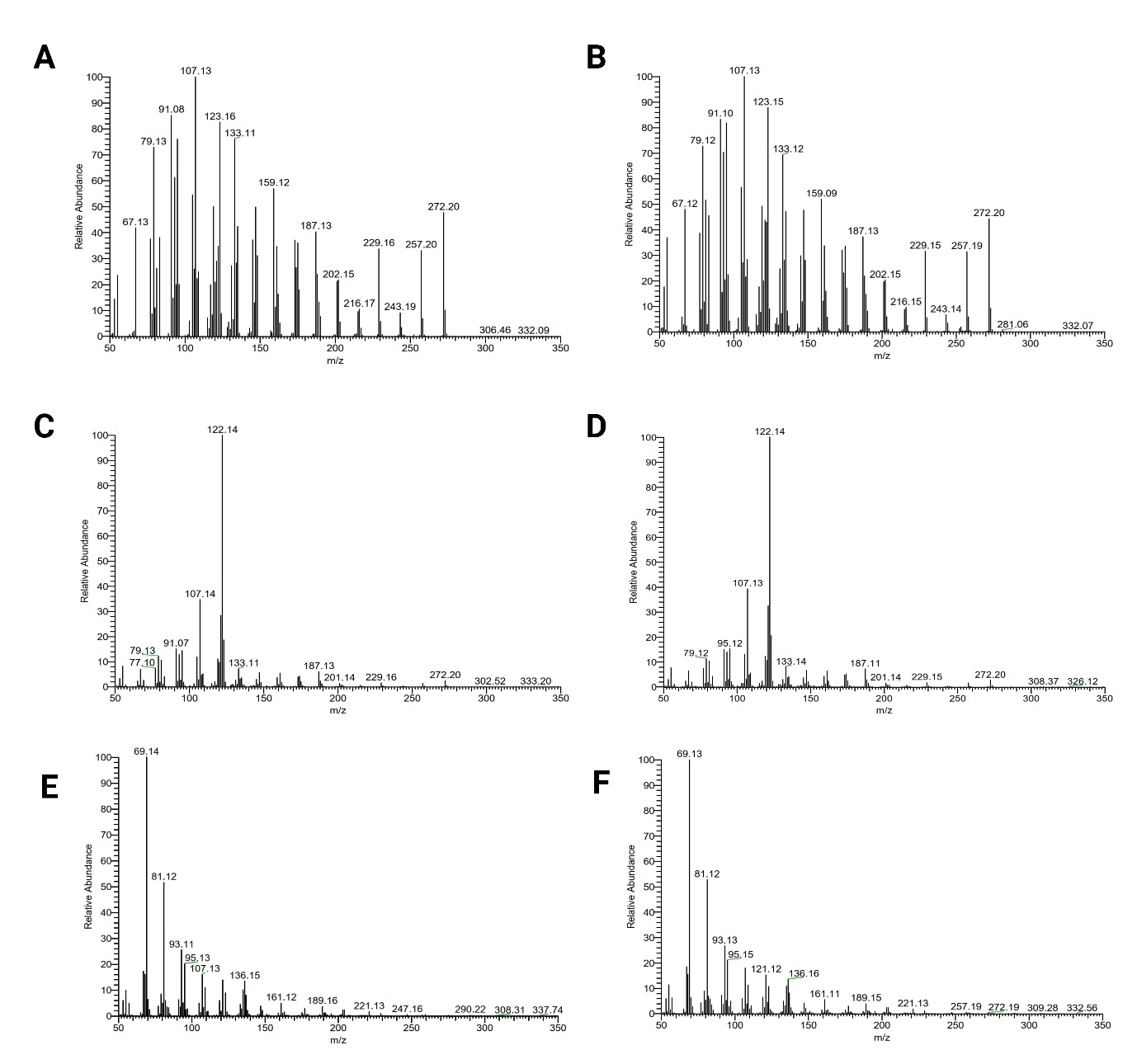

Figure S2. **Representative mass spectra of selected terpenes for LRS5 (A, C, E) and E_LRS5 (B, D, F).** A & B) iso-Taxadiene; C & D) Taxadiene; E & F) GGOH. Figure was created with BioRender.com.

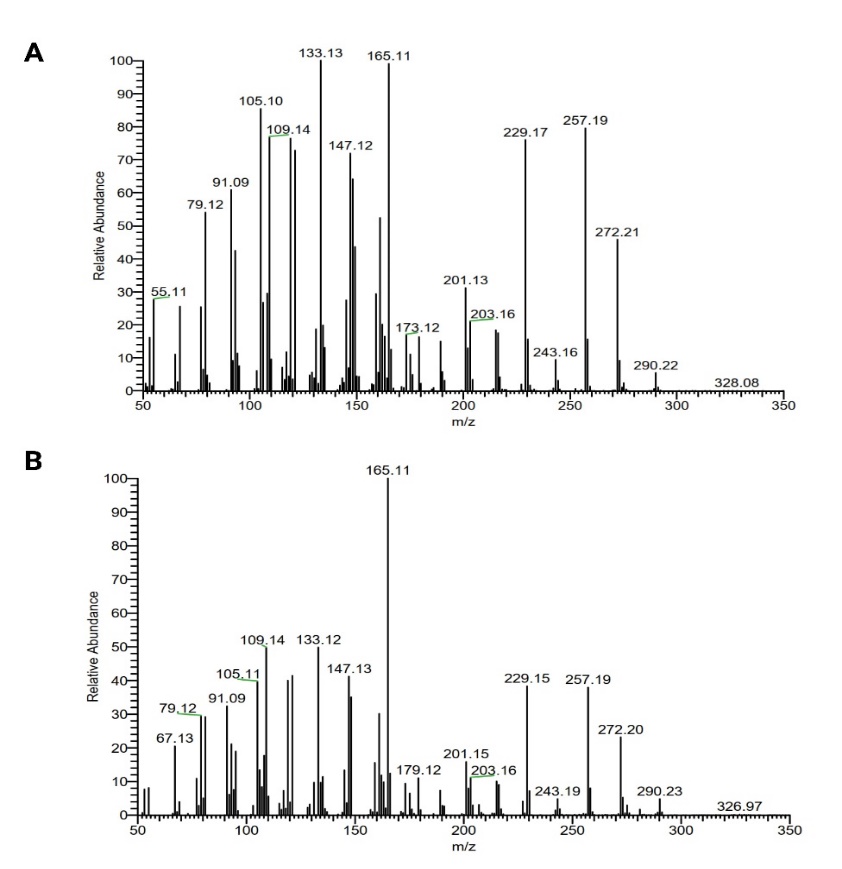

Figure S3. **Mass spectrum of an unknown terpenoid, eluted after taxadiene at 7:32 minutes.** A) LRS5; B) E_LRS5. Figure was created with BioRender.com.

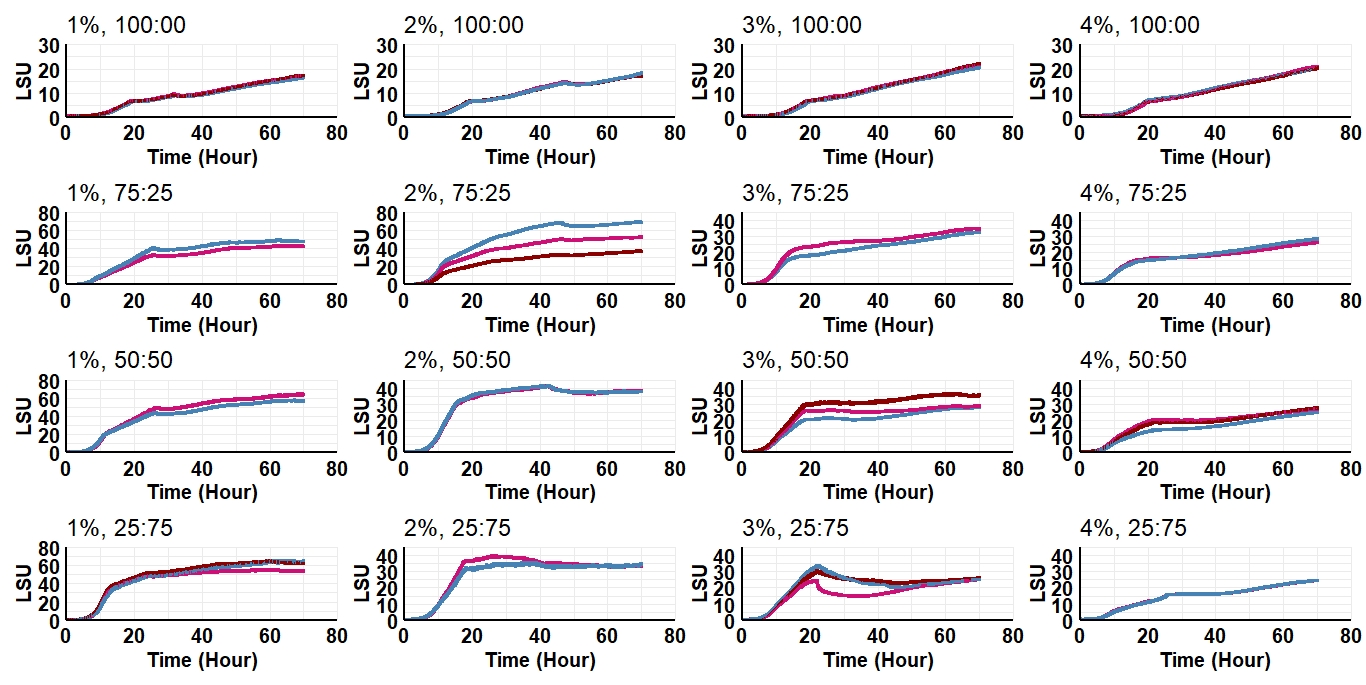

Figure S4. **Growth of LRS5 in buffered SD medium (pH = 6).** The cell was grown using 48-well FlowerPlate in BioLector, at a shaking speed of 1000 rpm and 30 °C incubation temperature. The SD medium contained different total sugar concentrations (1%-4%), at different galactose: glucose ratios of 100:00, 75:25, 50:50 and 25:75. The experiment was performed in triplicates. Any clear outlier was excluded from plotting and further analysis. LSU stands for light scattering unit.

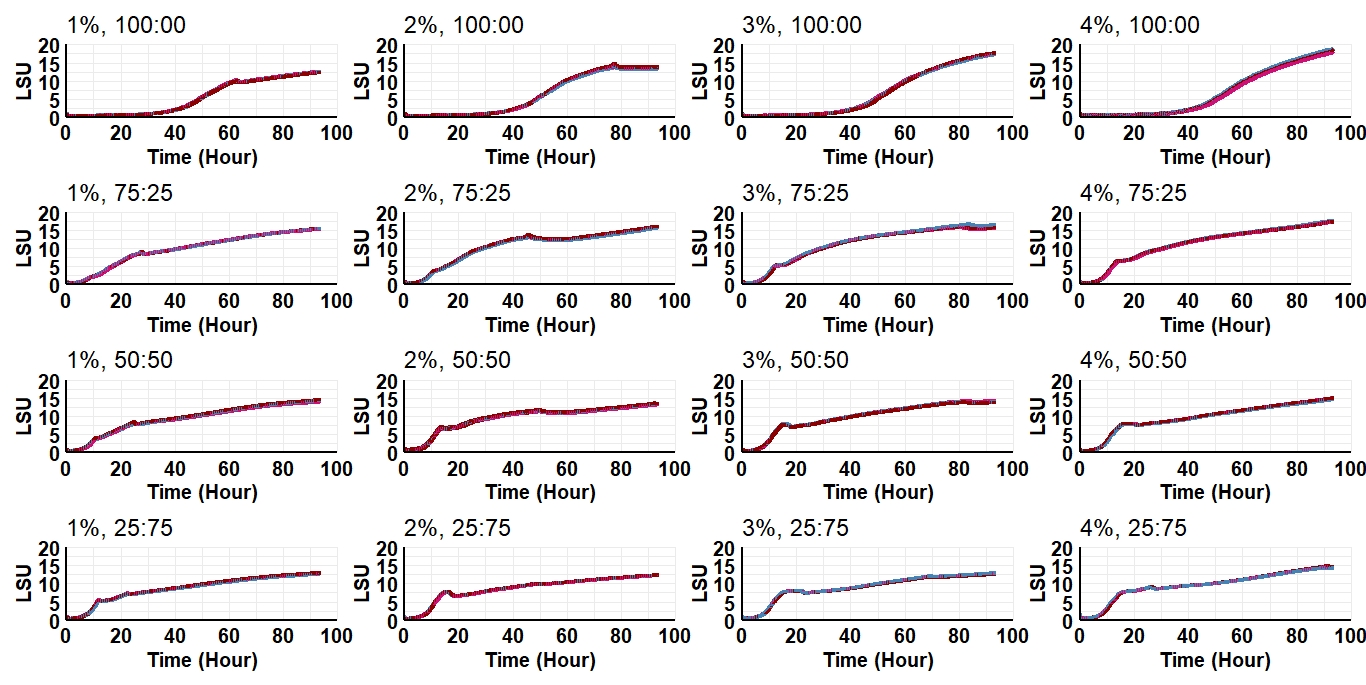

Figure S5. **Growth of E_LRS5 in buffered SD medium (pH = 6).** The cell was grown using 48-well FlowerPlate in BioLector, at a shaking speed of 1000 rpm and 30 °C incubation temperature. The SD medium contained different total sugar concentrations (1%-4%), at different galactose: glucose ratios of 100:00, 75:25, 50:50 and 25:75. The experiment was performed in triplicates. LSU stands for light scattering unit.

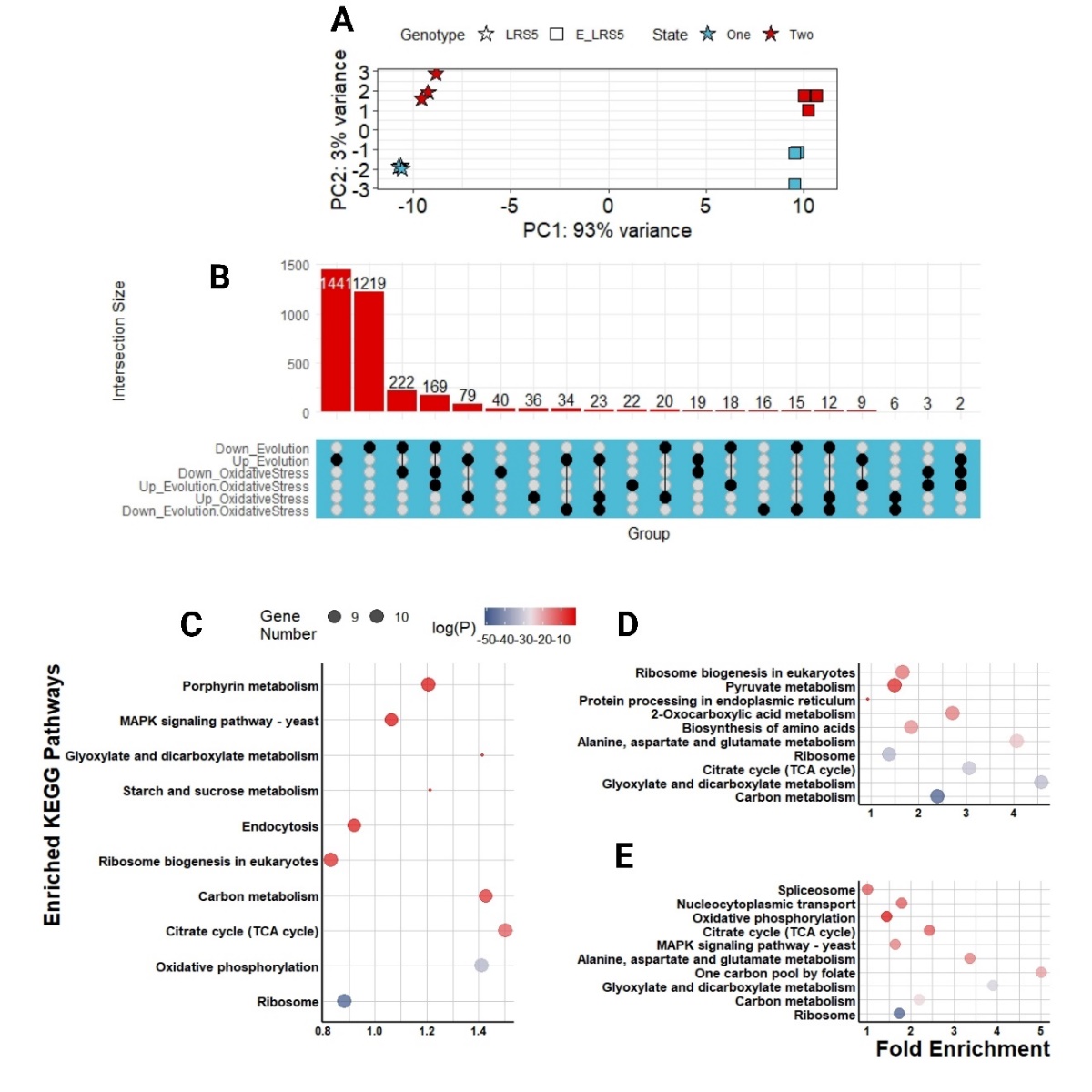

Figure S6. **RNA-seq and** **differential expression analysis.** A) Principal component analysis plot of samples. ″One″ and ″Two″ represent the steady state I (SSI) and steady state II (SSII) of continuous cultures for parent (LRS5) and evolved (E_LRS5) strains. B) Upset plot representing the contribution of each factor (Evolution and Oxidative Stress) and intersections of their contribution to the significant changes in gene expressions (Evolution. Oxidative Stress). Rows correspond to factors and the bar chart displays the number of single and common differentially expressed genes (DEGs) across all factors. Top ten enriched pathways, with the highest fold enrichment values, were plotted with respect to differentially expressed gene sets due to C) evolution, D) oxidative stress and E) evolution and oxidative stress effects (FDR cut-off < 0.05). The size of bubbles in the last three dot plots indicates the number of differentially expressed genes in each pathway. The colour bar shows the log10(P) values of the pathway, where shifting towards the blue colour shows more significant enrichment of the pathway. Figure was created with BioRender.com.

Table S1. **Physiological parameters for LRS5 and E_LRS5 in galactose-limited chemostats before (steady state I) and during (steady state II) H_2_O_2_ (0.5 mM) induction at 0.1 hr^-1^ dilution rate ^ab^**

| **Condition** | **Y**  **_Biomass_**  **_(g.g_^-1^_)_** | **q**  **_Galactose_** | **q/Y**  **_Acetate_** | **q/Y**  **_Ethanol_** | **q/Y**  **_Glycerol_** | **q/Y**  **_Pyruvate_** | **q/Y**  **_Succinate_** | **q/Y**  **_CO2_** | **Carbon Recovery** |
| --- | --- | --- | --- | --- | --- | --- | --- | --- | --- |
| LRS5_SSI | 0.27 ± 0.02 | 2.04 ± 0.19 | 0.**51** ± 0.0**2/**  0.066 ± 0.002 | **1.8** ± 0.**15/**  0.23 ± 0.019 | 0.077 ± 0.003/  0.015 ± 0.0012 | 0.025 ± 0.001/  0.005 ± 0.0003 | **0.000**3 ± 0.001**/**  8.08931E-05 ± 0.0001 | 5.38 ± 0.08/  0.349 ± 0.005 | 0.91 ± 0.05 |
| LRS5_SSII | 0.29 ± 0.01 | 1.88 ± 0.05 | 0.**5**2 ± 0.0**3**/  0.072 ± 0.004 | **1.79** ± 0.0**4/**  0.25 ± 0.006 | 0.056 ± 0.003/ 0.012 ± 0.0003 | 0.015 ± 0.002/ 0.003 ± 0.0004 | 0.00011± 0.0009/ 0.0001 ± 0.0002 | 5.09 ± 0.06/ 0.355 ± 0.004 | 0.96 ± 0.02 |
| E_LRS5_SSI | 0.34 ± 0.04 | 1.66 ± 0.18 | 0.**46** ± 0.0**6/**  0.067 ± 0.009 | **1.13** ± 0.0**8/**  0.165 ± 0.012 | 0.048 ± 0.01/ 0.010 ± 0.0005 | 0.042 ± 0.001/ 0.009 ± 0.0002 | 0/ 0 | 4.56 ± 0.2/ 0.333 ± 0.015 | 0.86 ± 0.05 |
| E_LRS5_SSII | 0.34 ± 0.01 | 1.65 ± 0.05 | 0.53 ± 0.0**4**/  0.077 ± 0.006 | **1.21** ± 0.0**9**/  0.17 ± 0.013 | 0.042 ± 0.002/ 0.009 ± 0.0006 | 0.037± 0.001/ 0.008 ± 0.0001 | 0/ 0 | 4.6 ± 0.25/ 0.329 ± 0.018 | 0.87 ± 0.05 |

^a^ Fluxes (q) are expressed as millimoles per gram of dry yeast biomass per hour.

^b^ Carbon recoveries/ yields (Y) were calculated based on yeast weight of CH_1.748_N_0.148_O_0.596_P_0.009_S_0.0019_M_0.018_, equal to 26.4 gDW/molC at dilution rate of 0.1 h^-1 1^. The unit is mmol C per mmol consumed galactose.

Table S2. **All enriched KEGG pathways and genes due to evolution effect.** The gene names are listed according to Ensembl gene nomenclature.

| **ID** | **Term_**  **Description** | **Fold_**  **Enrichment** | **lowest_p** | **highest_p** | **Up_**  **regulated** | **Down_**  **regulated** |
| --- | --- | --- | --- | --- | --- | --- |
| sce03010 | Ribosome | 0.883059 | 2.00E-51 | 5.10E-50 | YBR146W, YBR251W, YCR003W, YCR031C, YCR046C, YDL202W, YDR041W, YDR115W, YDR116C, YDR237W, YDR337W, YDR405W, YER050C, YGL068W, YGR027C, YGR220C, YHL004W, YHL033C, YHR147C, YJL063C, YJL096W, YJR113C, YKL003C, YML009C, YML025C, YMR121C, YMR188C, YMR193W, YMR286W, YNL081C, YNL185C, YNL284C, YNL306W, YNR036C, YOR150W, YPL013C, YPL173W, YPL183W-A, YPL249C-A, YPR166C | YBR031W, YDR012W, YDR025W, YFR031C-A, YIL052C, YIL133C, YJL191W, YKL180W, YLL045C, YLR009W, YLR029C, YLR048W, YLR061W, YLR075W, YLR167W, YLR185W, YLR264W, YLR325C, YLR333C, YLR340W, YLR344W, YLR367W, YLR388W, YLR406C, YLR441C, YLR448W, YML073C, YNL067W, YNL096C, YOL120C, YOR063W, YPL198W |
| sce00190 | Oxidative phosphorylation | 1.410127 | 9.30E-40 | 1.90E-34 | YBL045C, YBR011C, YBR039W, YCL005W-A, YDL004W, YDL067C, YDR178W, YDR298C, YDR322C-A, YDR377W, YDR529C, YEL024W, YEL051W, YER141W, YFR033C, YGL008C, YGL187C, YGL191W, YGR020C, YGR183C, YHR001W-A, YHR039C-A, YHR051W, YJL045W, YJL166W, YJR048W, YJR121W, YKL016C, YKL141W, YKL148C, YKL192C, YLL009C, YLL041C, YLR038C, YLR295C, YML081C-A, YML120C, YMR118C, YMR256C, YMR267W, YNL052W, YOL077W-A, YOR020W-A, YOR065W, YPL078C, YPL132W, YPL172C, YPL271W, YPR020W, YPR191W | Q0130, YDL085W, YDL185W, YEL039C, YKL080W, YLR164W, YLR447C, YMR145C, YOR270C |
| sce00020 | Citrate cycle (TCA cycle) | 1.503416 | 3.60E-16 | 1.00E-04 | YCR005C, YDR148C, YDR178W, YFL018C, YGR244C, YJL045W, YKL085W, YKL141W, YKL148C, YLL041C, YLR174W, YMR118C, YNL009W, YNL037C, YNL071W, YOR136W, YOR142W, YPL262W, YPR001W | YBR218C, YIL125W, YKR097W, YLR164W, YLR304C, YOL126C |
| sce01200 | Carbon metabolism | 1.427568 | 1.70E-09 | 3.90E-02 | YBR117C, YCL040W, YCL064C, YCR005C, YCR012W, YCR073W-A, YDL168W, YDR036C, YDR050C, YDR148C, YDR178W, YER081W, YER099C, YFL018C, YFL030W, YFR053C, YGR043C, YGR088W, YGR244C, YGR248W, YGR256W, YHL011C, YHR163W, YJL045W, YJL068C, YJL121C, YJR009C, YKL085W, YKL141W, YKL148C, YKL152C, YLL041C, YLR174W, YMR118C, YMR323W, YNL009W, YNL037C, YNL071W, YNL117W, YOL056W, YOR136W, YOR142W, YOR347C, YPL262W, YPR001W, YPR006C | YAL054C, YBR196C, YBR218C, YDR019C, YDR516C, YER065C, YER086W, YFL053W, YGL205W, YGL253W, YGR205W, YGR240C, YGR254W, YHR174W, YHR183W, YIL074C, YIL125W, YIR031C, YJL052W, YKL181W, YKR009C, YKR097W, YLR027C, YLR058C, YLR089C, YLR153C, YLR164W, YLR303W, YLR304C, YLR354C, YLR377C, YLR446W, YMR189W, YMR205C, YNL241C, YOL061W, YOL126C, YPL028W, YPR074C |
| sce03008 | Ribosome biogenesis in eukaryotes | 0.831349 | 2.90E-06 | 5.50E-05 | YAL033W, YCL031C, YCR057C, YDL166C, YGR276C, YHR148W, YHR196W, YOL010W, YOR039W, YOR119C, YOR185C | YGL173C, YGR090W, YGR218W, YIL035C, YJL109C, YLL034C, YLR022C, YLR106C, YLR107W, YLR129W, YLR145W, YLR175W, YLR186W, YLR197W, YLR222C, YLR293C, YLR397C, YLR409C, YNL163C, YNR053C, YPL169C, YPL217C |
| sce00630 | Glyoxylate and dicarboxylate metabolism | 1.414248 | 1.70E-03 | 1.70E-03 | YCR005C, YFL018C, YFL030W, YGR088W, YKL085W, YNL117W, YNL274C, YPR001W, YPR006C | YAL054C, YBR222C, YDR019C, YER065C, YGR205W, YIR031C, YLR058C, YLR153C, YLR304C, YMR189W, YOL126C, YPL028W, YPR035W |
| sce04144 | Endocytosis | 0.920319 | 1.80E-03 | 3.10E-03 | YBL075C, YBR234C, YCL008C, YDL137W, YDL192W, YER031C, YFL005W, YGR206W, YKL002W, YML001W, YOR357C, YPL065W | YAL005C, YBL047C, YDL229W, YDR153C, YDR170C, YDR524C, YER125W, YER144C, YGL095C, YGL206C, YHL002W, YIL034C, YIL044C, YIL062C, YJL154C, YJR005W, YKL079W, YLL024C, YLR025W, YLR181C, YLR206W, YLR229C, YLR370C, YLR417W, YNL209W, YNR006W, YOR069W |
| sce04011 | MAPK signaling pathway - yeast | 1.062942 | 2.40E-03 | 2.30E-02 | YBR083W, YCL027W, YCL032W, YCR073C, YCR084C, YDL006W, YDL022W, YDR144C, YDR480W, YDR490C, YGR070W, YGR088W, YIR019C, YJL128C, YMR036C, YNL283C, YOR231W, YPL187W, YPR119W, YPR120C | YAL041W, YBL105C, YDR103W, YDR379W, YDR420W, YER125W, YER155C, YGR014W, YGR032W, YHR030C, YIL015W, YIL113W, YIL147C, YIL159W, YKL101W, YKL161C, YLL021W, YLR006C, YLR079W, YLR113W, YLR120C, YLR182W, YLR229C, YLR305C, YLR332W, YLR342W, YLR362W, YLR371W, YLR425W, YMR037C, YMR043W, YMR164C, YMR199W, YNL271C, YOL059W, YOL105C, YOL113W, YOL151W, YOR008C, YOR127W, YOR208W, YPL049C, YPL089C, YPL256C, YPR075C |
| sce00860 | Porphyrin metabolism | 1.20627 | 3.20E-03 | 4.40E-03 | YDL120W, YDL205C, YDR047W, YER141W, YGL040C, YKL087C, YKR069W, YOL033W, YOR278W, YPL172C | YAL039C |
| sce04070 | Phosphatidylinositol signaling system | 1.053699 | 8.80E-03 | 8.80E-03 | YDR173C | YBL105C, YFR019W, YHR046C, YIL002C, YKL212W, YLR240W, YLR305C, YLR410W, YNL106C, YNL267W, YOR109W, YPR113W |
| sce00051 | Fructose and mannose metabolism | 1.610022 | 9.50E-03 | 1.20E-02 | YCL040W, YDR050C, YER003C, YFR053C | YDL055C, YDL246C, YDR516C, YFL053W, YGL253W, YGR240C, YIL107C, YJL155C, YJR159W, YLR345W, YLR377C, YLR446W, YMR205C, YNR073C, YOL136C |
| sce00500 | starch and sucrose metabolism | 1.211754 | 9.90E-03 | 1.10E-02 | YBR126C, YBR299W, YCL040W, YDR074W, YEL011W, YFR015C, YFR053C, YGR287C, YGR292W, YHL012W, YJL137C, YMR105C, YOR190W, YPR160W | YBR196C, YDR516C, YGL253W, YGR032W, YGR282C, YIL099W, YIL162W, YKL127W, YLR258W, YLR300W, YLR342W, YLR446W |
| sce00620 | Pyruvate metabolism | 1.242824 | 2.80E-02 | 2.80E-02 | YBL015W, YDL168W, YDL178W, YDR533C, YFL018C, YGL256W, YKL085W, YML004C, YML054C, YMR169C, YNL071W, YNL117W, YOR108W, YOR347C, YPL262W | YAL054C, YBR218C, YDL131W, YDL174C, YDL182W, YEL071W, YER073W, YIR031C, YKR097W, YLR153C, YMR110C, YMR303C, YNL104C, YNR016C, YOL126C, YOL151W, YPL028W, YPL061W, YPL280W |

Table S3. **All enriched KEGG pathways and genes due to oxidative stress effect.** The gene names are listed according to Ensembl gene nomenclature.

| **ID** | **Term_**  **Description** | **Fold_**  **Enrichment** | **Lowest_p** | **highest_p** | **Up_**  **regulated** | **Down_**  **regulated** |
| --- | --- | --- | --- | --- | --- | --- |
| sce01200 | Carbon metabolism | 2.401316 | 3.20E-23 | 2.80E-17 | YAL044C, YDR019C, YER065C, YFL030W, YGL205W, YGR043C, YKL085W, YKR009C, YKR097W, YLR174W, YLR377C, YMR189W, YNL037C, YNL117W, YOR136W, YPR006C | YGL062W, YJL045W, YLL041C, YLR058C, YLR089C, YLR153C, YLR304C, YLR354C, YLR446W, YOL126C, YPL028W, YPR074C |
| sce00630 | Glyoxylate and dicarboxylate metabolism | 4.595622 | 1.00E-18 | 1.80E-16 | YAL044C, YDR019C, YER065C, YFL030W, YKL085W, YMR189W, YNL117W, YPR006C | YLR058C, YLR153C, YLR304C, YOL126C, YPL028W, YPR035W |
| sce00020 | Citrate cycle (TCA cycle) | 3.070808 | 3.70E-16 | 8.40E-12 | YKL085W, YKR097W, YLR174W, YNL037C, YOR136W | YGL062W, YJL045W, YLL041C, YLR304C, YOL126C |
| sce03010 | Ribosome | 1.377823 | 3.20E-15 | 5.60E-14 | YFL034C-A, YHL033C, YKL003C | YDL082W, YDL130W, YDR382W, YFR031C-A, YLL045C, YLR009W, YLR048W, YLR061W, YLR167W, YLR185W, YLR325C, YLR333C, YLR340W, YLR344W, YLR406C, YLR441C, YLR448W, YNL069C, YNL096C |
| sce00250 | Alanine, aspartate and glutamate metabolism | 4.079787 | 4.00E-13 | 1.30E-10 | YAL062W, YDL215C, YFL030W | YDL171C, YHR018C, YKL104C, YLR089C, YLR351C, YLR359W, YOL058W, YPR035W, YPR145W |
| sce01230 | Biosynthesis of amino acids | 1.842485 | 8.50E-12 | 3.40E-08 | YGR043C, YLR174W, YNL037C, YOR136W | YBR249C, YDL171C, YDR035W, YDR234W, YGL009C, YGL062W, YHR018C, YJL200C, YJR016C, YLL058W, YLR058C, YLR089C, YLR180W, YLR304C, YLR354C, YLR355C, YOL058W, YPR035W, YPR074C, YPR145W |
| sce01210 | 2-Oxocarboxylic acid metabolism | 2.719858 | 6.00E-11 | 1.20E-07 | YLR174W, YNL037C, YOR136W | YDR234W, YGL009C, YJL200C, YJR016C, YLR089C, YLR304C, YLR355C |
| sce00190 | Oxidative phosphorylation | 1.220449 | 3.60E-10 | 5.20E-06 | YKL016C, YKL192C, YMR054W, YPL172C | YEL024W, YJL045W, YJR048W, YLL041C, YLR395C, YLR447C |
| sce04141 | Protein processing in endoplasmic reticulum | 0.921242 | 6.70E-10 | 5.00E-03 | YOL013C | YLL024C, YLR057W, YLR080W, YLR090W, YLR207W, YLR208W, YLR378C, YML130C |
| sce03008 | Ribosome biogenesis in eukaryotes | 1.672345 | 3.20E-09 | 5.70E-09 |  | YLL034C, YLR022C, YLR059C, YLR106C, YLR129W, YLR145W, YLR175W, YLR186W, YLR197W, YLR222C, YLR293C, YLR397C, YLR409C |
| sce00100 | steroid biosynthesis | 3.173168 | 1.70E-08 | 4.30E-08 |  | YGR175C, YHR007C, YHR072W, YLR100W, YML008C, YMR015C |
| sce04146 | Peroxisome | 2.141888 | 1.80E-08 | 4.40E-02 | YDR265W, YFL030W, YGL205W, YIL160C, YLR174W, YOL147C | YLR191W, YMR208W, YOR317W |
| sce04011 | MAPK signaling pathway - yeast | 1.252566 | 3.40E-08 | 1.30E-03 |  | YHR030C, YKL161C, YLL021W, YLR006C, YLR079W, YLR113W, YLR120C, YLR182W, YLR229C, YLR305C, YLR332W, YLR342W, YLR362W, YLR371W, YLR425W |
| sce00220 | Arginine biosynthesis | 3.919796 | 1.30E-07 | 1.60E-03 | YAL062W, YDL215C | YBR208C, YHR018C, YLR089C, YOL058W, YPR035W |
| sce00260 | Glycine, serine and threonine metabolism | 1.586584 | 2.80E-07 | 7.20E-05 | YAL044C, YDR019C, YFL030W, YMR189W | YLR058C |
| sce00030 | Pentose phosphate pathway | 1.359929 | 4.80E-07 | 3.90E-05 | YGR043C, YLR377C | YLR354C, YPR074C |
| sce00670 | One carbon pool by folate | 2.538534 | 3.70E-06 | 4.70E-04 | YDR019C, YKR080W | YLR028C, YLR058C |
| sce04111 | Cell cycle - yeast | 0.659043 | 3.70E-06 | 2.50E-05 |  | YLL004W, YLR079W, YLR086W, YLR102C, YLR127C, YLR182W, YLR272C, YLR274W, YLR288C |
| sce00071 | Fatty acid degradation | 2.379876 | 6.40E-06 | 1.80E-03 | YGL205W, YIL160C, YMR303C | YOR317W, YPL028W |
| sce01212 | Fatty acid metabolism | 2.596228 | 9.60E-06 | 1.40E-05 | YGL055W, YGL205W, YIL160C | YLR372W, YOR317W, YPL028W |
| sce00620 | Pyruvate metabolism | 1.493256 | 1.10E-05 | 6.30E-04 | YKL085W, YKR097W, YMR303C, YNL117W | YGL062W, YLR153C, YOL126C, YPL028W |
| sce00051 | Fructose and mannose metabolism | 1.730819 | 1.40E-05 | 1.40E-05 | YLR377C | YER003C, YLR345W, YLR446W |
| sce04113 | Meiosis - yeast | 0.899481 | 9.70E-05 | 7.10E-04 | YDR342C, YDR343C, YHR096C | YLL004W, YLR079W, YLR081W, YLR102C, YLR127C, YLR182W, YLR263W, YLR274W, YLR288C |
| sce00640 | Propanoate metabolism | 2.379876 | 1.70E-04 | 3.90E-02 | YGL205W, YKR009C | YLR153C |
| sce03013 | Nucleocytoplasmic transport | 1.208826 | 3.10E-04 | 3.10E-04 | YNL139C | YLL023C, YLR018C, YLR064W, YLR208W, YLR293C, YLR335W, YLR347C |
| sce04139 | Mitophagy - yeast | 1.427926 | 4.20E-04 | 1.90E-02 |  | YHR030C, YLL001W, YLL006W, YLR006C, YLR113W, YLR356W |
| sce00410 | beta-Alanine metabolism | 1.464539 | 4.70E-04 | 4.60E-02 | YGL205W, YKR009C | |
| sce00010 | Glycolysis/Gluconeogenesis | 1.057723 | 5.70E-04 | 1.10E-03 | YKR097W, YLR377C, YMR303C | YLR134W, YLR153C, YLR446W |
| sce04130 | sNARE interactions in vesicular transport | 1.903901 | 1.20E-03 | 1.50E-03 | YOL018C | YLR026C, YLR078C, YLR268W |
| sce00900 | Terpenoid backbone biosynthesis | 1.427926 | 1.20E-03 | 1.50E-03 |  | YLR450W, YMR208W, YPL028W |
| sce00230 | Purine metabolism | 1.23401 | 1.40E-03 | 1.40E-03 | YER070W, YKL067W | YCL050C, YLR028C, YLR209C, YLR359W, YLR432W |
| sce01250 | Biosynthesis of nucleotide sugars | 1.241674 | 1.80E-03 | 1.80E-03 |  | YER003C, YKL104C, YLR446W |
| sce03018 | RNA degradation | 0.906619 | 2.70E-03 | 4.80E-02 |  | YLR187W, YLR259C, YLR270W, YLR369W, YLR398C, YLR438C-A |
| sce00480 | Glutathione metabolism | 2.069457 | 2.80E-03 | 2.80E-03 | YER070W, YLR174W | YLR146C, YLR299W, YPL091W |
| sce00240 | Pyrimidine metabolism | 1.313035 | 3.70E-03 | 3.70E-03 | YER070W, YKL067W | YLR245C, YLR420W |
| sce00520 | Amino sugar and nucleotide sugar metabolism | 1.730819 | 5.50E-03 | 5.50E-03 | YKL150W | YER003C, YKL104C, YLR286C, YLR307W, YLR446W |
| sce01232 | Nucleotide metabolism | 1.427926 | 9.80E-03 | 9.80E-03 | YER070W, YKL067W | YLR209C, YLR245C, YLR359W, YLR432W |
| sce00500 | starch and sucrose metabolism | 1.189938 | 9.80E-03 | 9.80E-03 |  | YDR261C, YLR258W, YLR300W, YLR342W, YLR446W |
| sce01040 | Biosynthesis of unsaturated fatty acids | 3.807801 | 1.90E-02 | 2.60E-02 | YGL055W, YGL205W, YIL160C | YLR372W |
| sce00300 | Lysine biosynthesis | 1.586584 | 3.90E-02 | 3.90E-02 |  | YDR234W, YJL200C |
| sce00280 | Valine, leucine and isoleucine degradation | 1.359929 | 3.90E-02 | 4.60E-02 | YIL160C | YPL028W |

Table S4. **All enriched KEGG pathways and genes due to evolution and oxidative stress interaction effect.** The gene names are listed according to Ensembl gene nomenclature.

| **ID** | **Term_**  **Description** | **Fold_**  **Enrichment** | **lowest_**  **p** | **highest_**  **p** | **Up_**  **regulated** | **Down_**  **regulated** |
| --- | --- | --- | --- | --- | --- | --- |
| sce03010 | Ribosome | 1.735134 | 1.10E-16 | 3.00E-16 | YBR191W, YIL052C, YLL045C, YLR048W, YLR075W, YLR167W, YLR185W, YLR325C, YLR333C, YLR344W, YLR406C, YLR441C, YLR448W | YHL033C |
| sce01200 | Carbon metabolism | 2.206322 | 6.40E-12 | 9.90E-10 | YLR153C | YDR019C, YER065C, YFL030W, YGL205W, YKR009C, YKR097W, YLR377C, YMR189W, YNL037C, YNL117W, YOR136W, YPL262W |
| sce00630 | Glyoxylate and dicarboxylate metabolism | 3.897641 | 3.70E-11 | 6.30E-09 | YLR153C | YDR019C, YER065C, YFL030W, YMR189W, YNL117W |
| sce00670 | One carbon pool by folate | 5.023626 | 9.50E-10 | 4.60E-07 | YLR028C | YDR019C, YKR080W, YMR120C |
| sce04011 | MAPK signaling pathway - yeast | 1.652509 | 8.20E-09 | 2.00E-07 | YHR030C, YLR006C, YLR113W, YLR120C, YLR182W, YLR229C, YLR332W, YLR362W, YLR371W | YJL157C |
| sce00250 | Alanine, aspartate and glutamate metabolism | 3.364035 | 1.90E-07 | 3.80E-06 | YLR351C | YAL062W, YDL215C, YFL030W, YJR109C |
| sce00020 | Citrate cycle (TCA cycle) | 2.430787 | 3.30E-07 | 1.50E-04 |  | YKR097W, YNL037C, YOR136W, YPL262W |
| sce00190 | Oxidative phosphorylation | 1.449123 | 1.10E-06 | 3.10E-02 | YLR295C, YLR395C, YLR447C, YOR020W-A | YGL008C, YJR121W |
| sce03013 | Nucleocytoplasmic transport | 1.794152 | 2.20E-06 | 2.20E-06 | YLR018C, YLR064W, YLR208W, YLR293C, YLR335W | YKR095W |
| sce03040 | spliceosome | 1.004725 | 2.40E-06 | 2.40E-06 | YBR065C, YLL036C, YLR147C, YLR298C | |
| sce00260 | Glycine, serine and threonine metabolism | 1.88386 | 9.20E-06 | 8.10E-04 |  | YDR019C, YFL030W, YMR189W |
| sce00620 | Pyruvate metabolism | 1.846921 | 2.30E-05 | 2.30E-05 | YLR153C | YKR097W, YMR303C, YNL117W, YPL262W |
| sce00640 | Propanoate metabolism | 4.709649 | 2.80E-05 | 2.80E-05 | YLR153C | YGL205W, YKR009C |
| sce04141 | Protein processing in endoplasmic reticulum | 1.215393 | 3.10E-05 | 5.30E-04 | YBR082C, YBR101C, YLR080W, YLR090W, YLR208W, YML130C | |
| sce00410 | beta-Alanine metabolism | 2.898246 | 3.60E-05 | 3.60E-05 |  | YGL205W, YKR009C |
| sce00920 | sulfur metabolism | 2.511813 | 5.80E-05 | 5.80E-05 | YLL058W | YJR137C |
| sce00230 | Purine metabolism | 1.395452 | 5.80E-05 | 5.80E-05 | YBR111C, YLR028C | YGR061C, YMR120C |
| sce00100 | steroid biosynthesis | 3.139766 | 1.00E-04 | 1.00E-04 | YHR072W, YLR056W, YLR100W | |
| sce00220 | Arginine biosynthesis | 2.216306 | 1.40E-04 | 3.20E-02 |  | YAL062W, YDL215C |
| sce00071 | Fatty acid degradation | 2.82579 | 1.40E-04 | 1.40E-04 |  | YGL205W, YMR246W, YMR303C |
| sce04139 | Mitophagy - yeast | 2.354825 | 2.40E-04 | 5.70E-03 | YHR030C, YLR006C, YLR113W, YLR147C, YLR356W | |
| sce03050 | Proteasome | 1.046589 | 8.90E-04 | 8.90E-04 | YBR173C, YLR421C | |
| sce01210 | 2-Oxocarboxylic acid metabolism | 1.076491 | 1.30E-03 | 1.30E-02 |  | YNL037C, YOR136W |
| sce04146 | Peroxisome | 1.412895 | 2.00E-03 | 1.90E-02 |  | YFL030W, YGL205W, YMR246W |
| sce00010 | Glycolysis/Gluconeogenesis | 1.395452 | 3.10E-03 | 3.10E-03 | YLR153C | YKR097W, YLR377C, YMR303C |
| sce04120 | Ubiquitin mediated proteolysis | 1.88386 | 3.90E-03 | 3.90E-03 | YBR082C, YLL036C, YLR102C, YLR127C, YLR167W | |
| sce04111 | Cell cycle - yeast | 0.724561 | 5.80E-03 | 5.80E-03 | YLR102C, YLR127C, YLR182W, YLR288C | YJL157C |
| sce01040 | Biosynthesis of unsaturated fatty acids | 3.767719 | 6.10E-03 | 6.10E-03 | YLR372W | YGL205W |
| sce04144 | Endocytosis | 1.43078 | 2.30E-02 | 2.30E-02 | YLR025W, YLR206W, YLR229C, YLR370C, YLR417W | YKR031C |
| sce03250 | Viral life cycle - HIV-1 | 2.511813 | 3.00E-02 | 3.70E-02 | YLR025W, YLR293C | |
| sce01212 | Fatty acid metabolism | 3.425199 | 3.10E-02 | 3.10E-02 | YLR372W | YGL205W, YKL182W, YMR246W |
| sce00380 | Tryptophan metabolism | 0.94193 | 3.40E-02 | 3.40E-02 | YLR231C |  |
| sce01230 | Biosynthesis of amino acids | 0.455773 | 3.70E-02 | 3.70E-02 | YLL058W | YNL037C, YOR136W |
| sce04070 | Phosphatidylinositol signalling system | 0.819069 | 4.50E-02 | 4.50E-02 | YBR109C |  |
